# Multilayer meta-matching: translating phenotypic prediction models from multiple datasets to small data

**DOI:** 10.1101/2023.12.05.569848

**Authors:** Pansheng Chen, Lijun An, Naren Wulan, Chen Zhang, Shaoshi Zhang, Leon Qi Rong Ooi, Ru Kong, Jianzhong Chen, Jianxiao Wu, Sidhant Chopra, Danilo Bzdok, Simon B Eickhoff, Avram J Holmes, B.T. Thomas Yeo

## Abstract

Resting-state functional connectivity (RSFC) is widely used to predict phenotypic traits in individuals. Large sample sizes can significantly improve prediction accuracies. However, for studies of certain clinical populations or focused neuroscience inquiries, small-scale datasets often remain a necessity. We have previously proposed a “meta-matching” approach to translate prediction models from large datasets to predict new phenotypes in small datasets. We demonstrated large improvement of meta-matching over classical kernel ridge regression (KRR) when translating models from a single source dataset (UK Biobank) to the Human Connectome Project Young Adults (HCP-YA) dataset. In the current study, we propose two meta-matching variants (“meta-matching with dataset stacking” and “multilayer meta-matching”) to translate models from multiple source datasets across disparate sample sizes to predict new phenotypes in small target datasets. We evaluate both approaches by translating models trained from five source datasets (with sample sizes ranging from 862 participants to 36,834 participants) to predict phenotypes in the HCP-YA and HCP-Aging datasets. We find that multilayer meta-matching modestly outperforms meta-matching with dataset stacking. Both meta-matching variants perform better than the original “meta-matching with stacking” approach trained only on the UK Biobank. All meta-matching variants outperform classical KRR and transfer learning by a large margin. In fact, KRR is better than classical transfer learning when less than 50 participants are available for finetuning, suggesting the difficulty of classical transfer learning in the very small sample regime. The multilayer meta-matching model is publicly available at GITHUB_LINK.

## 1. Introduction

There is growing interest in harnessing neuroimaging data to predict non-neuroimaging-related phenotypes, such as fluid intelligence or clinical outcomes, of individual participants (Gabrieli et al., 2015; Woo et al., 2017; Eickhoff & Langner, 2019; Varoquaux & Poldrack, 2019). However, most brain-behavior prediction studies suffer from underpowered samples, typically involving less than a few hundred participants, leading to low reproducibility and inflated performance (Arbabshirani et al., 2017; Bzdok & Meyer-Lindenberg, 2018; Masouleh et al., 2019; Poldrack et al., 2020; Marek et al., 2022). Adequately powered sample sizes can significantly improve prediction accuracy (Chu et al., 2012; Cui & Gong, 2018; He et al., 2020; Schulz et al., 2020), so large-scale datasets, such as the UK Biobank (Sudlow et al., 2015; Miller et al., 2016), are vital for enhancing prediction performance. However, for investigations of certain clinical populations or focused neuroscience inquiries, small-scale datasets often remain the norm.

We have previously proposed a “meta-matching” approach to translate prediction models from large datasets to improve the prediction of new phenotypes in small datasets (He et al., 2022). Meta-matching is grounded in the observation that many phenotypes exhibit inter-correlations, as demonstrated by previous studies identifying a small number of factors linking brain imaging data to various non-brain-imaging traits like cognition, mental health, demographics, and other health attributes (Smith et al., 2015; Miller et al., 2016; Xia et al., 2018; Kebets et al., 2019). As a result, a phenotype X in a smaller-scale study is likely correlated with a phenotype Y present in a larger population dataset. This means that a machine learning model trained on phenotype Y from the larger dataset might be more effectively translated to predict phenotype X in the smaller study. Meta-matching exploited these inter-phenotype correlations and was thus referred to as “meta-matching” given its close links with meta-learning (Fei-Fei et al., 2006; Andrychowicz et al., 2016; Finn et al., 2017; Ravi & Larochelle, 2016; Vanschoren, 2019). We note that meta-learning is also referred to “learning to learn” and is closely related to “transfer learning” (Hospedales et al., 2021). One distinction between meta-learning and transfer learning is that in transfer learning, the prediction problem in the target dataset can be same (Vakli et al., 2018; C.-L. Chen et al., 2020; Zhang & Bellec, 2020) or different (Hon & Khan, 2017; Lu et al., 2021; Schirmer et al., 2021) from the source dataset. On the other hand, meta-learning always involves training a machine learning model from a wide range of meta-training tasks and then adapting to perform a new prediction problem in the target dataset.

In our previous study (He et al., 2022), we trained a deep neural network (DNN) to predict 67 non-brain-imaging phenotypes from resting-state functional connectivity (RSFC) in the UK Biobank. The DNN was then translated using meta-matching to predict non-brain-imaging phenotypes in the Human Connectome Project Young Adult (HCP-YA) dataset, yielding large improvements over classical KRR without meta-learning. Among the different meta-matching variants, complementing basic meta-matching with stacking (which we will refer to as “meta-matching with stacking”) performed the best (He et al., 2022). Stacking is a well-known ensemble learning approach (Wolpert, 1992; Breiman, 1996) and has also enjoyed utility in neuroimaging (Liem et al., 2017; Rahim et al., 2017; Ooi et al., 2022).

The original study (He et al., 2022) experimented with only one source dataset (UK Biobank). Using multiple source datasets might lead to better generalization for multiple reasons. First, prediction performance tends to increase with larger sample sizes (Chu et al., 2012; Cui & Gong, 2018; He et al., 2020; Schulz et al., 2020). Second, given acquisition, preprocessing and demographic differences across datasets, training on multiple source datasets might yield representations that are more generalizable to a new target population (Abraham et al., 2017). Third, different datasets collect overlapping and distinct non-brain-imaging phenotypes. Since meta-matching exploits inter-phenotype correlation, training on more diverse phenotypes might lead to better performance. Here, we investigated the performance of meta-matching models trained from five source datasets - UK Biobank (Sudlow et al., 2015; Miller et al., 2016), Adolescent Brain Cognitive Development (ABCD) study (Volkow et al., 2018), Genomics Superstruct Project (GSP; Holmes et al., 2015), Healthy Brain Network (HBN; Alexander et al., 2017), and the enhanced Nathan Kline Institute-Rockland sample (eNKI-RS; Nooner et al., 2012).

One major challenge is the extreme sample size imbalances across datasets, e.g., the UK Biobank is almost 40 times larger than the HBN dataset. A second challenge is that the available phenotypes are different across datasets, so training a single DNN to predict all phenotypes is not straightforward. Here, we considered a naive extension of the original meta-matching with stacking approach by training independent prediction model(s) in each source dataset, and then performed stacking on the outputs of the prediction models in the target dataset. We refer to this extension as “meta-matching with dataset stacking”. Because meta-matching can improve the prediction of smaller datasets, we also proposed an alternative “multilayer meta-matching” approach, which gradually applied meta-matching from large source datasets (e.g., UK Biobank) to smaller source datasets (e.g., GSP, HBN, etc), to generate additional features for a final round of stacking in the target dataset.

We evaluated the proposed approaches in two target datasets - HCP-YA (Van Essen et al., 2013) and HCP-Aging (Harms et al., 2018). We found that both approaches performed better than the original “meta-matching with stacking” approach trained only on the UK Biobank. Given the close relationship between meta-learning and transfer learning, instead of performing stacking on the DNN trained on the UK Biobank (i.e., meta-matching with stacking), we also considered a standard transfer learning baseline (Weiss et al., 2016), in which the DNN was finetuned on the target dataset. Of note, meta-matching with stacking significantly outperformed the transfer learning baseline. In fact, the transfer learning baseline was worse than classical kernel ridge regression when less than 50 participants were available for finetuning, suggesting the difficulty of transfer learning in the very small sample regime. Finally, we found that multilayer meta-matching modestly outperformed meta-matching with dataset stacking.

## 2. Methods

### 2.1 Datasets

As illustrated in Figure 1, we used five source datasets for meta-training: the UK Biobank (Sudlow et al., 2015; Miller et al., 2016), the Adolescent Brain Cognitive Development (ABCD) study (Volkow et al., 2018), the Genomics Superstruct Project (GSP; Holmes et al., 2015), the Healthy Brain Network (HBN; Alexander et al., 2017) project, and the enhanced Nathan Kline Institute-Rockland sample (eNKI-RS; Nooner et al., 2012). The models from the five datasets were then adapted for phenotypic prediction in two meta-test datasets: Human Connectome Project Young Adults (HCP-YA; Van Essen et al., 2013) and HCP-Aging (Harms et al., 2018). All data collection and analysis procedures were approved by the respective Institutional Review Boards (IRBs), including the National University of Singapore IRB for the analysis presented in this paper.

**Figure 1.**
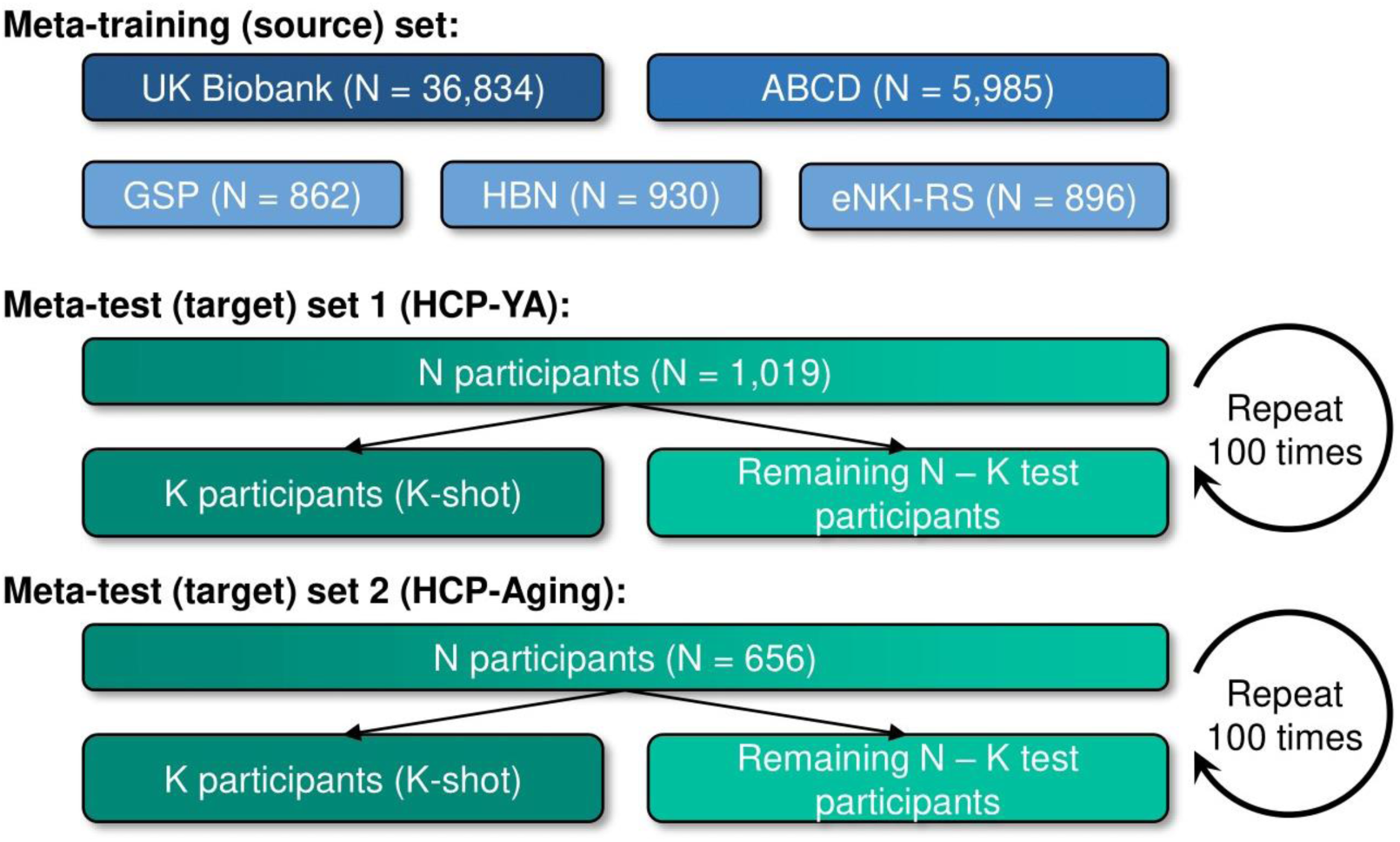
Schematic of meta-training and meta-test sets. Datasets were assigned to meta-training set and meta-test set. Prediction models from the meta-training set were adapted to K participants from each meta-test dataset to predict target phenotypes. The adapted models were evaluated in the remaining N – K participants from the meta-test dataset. This procedure was repeated 100 times for stability. The meta-training set was differentiated into extra-large-scale (UK Biobank; dark blue), large-scale (ABCD; blue) and medium-scale (GSP, HBN and eNKI-RS; light blue) source datasets.

The summary information of the datasets is listed in Table 1. Detailed information about the non-brain-imaging phenotypes (henceforth referred to as phenotypes) used can be found in Tables S2 to S8. The phenotypes covered a broad range of behavioral domains, ranging from cognitive performance, personality measures, lifestyle and mental health scores. The following subsections describe each dataset and corresponding preprocessing procedures in greater detail.

**Table 1.**
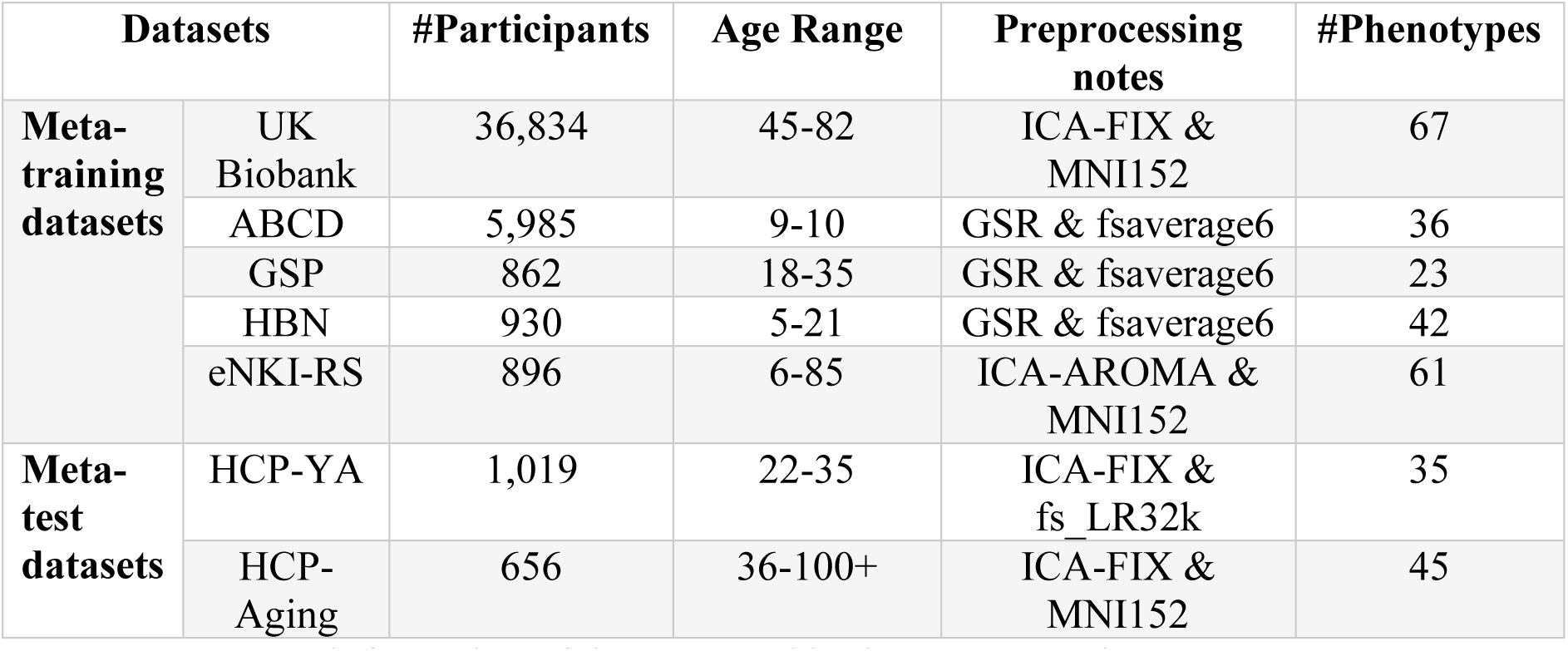
Summary information of datasets used in the current study.

| Datasets |  | #Participants | Age Range | Preprocessing notes | #Phenotypes |
| --- | --- | --- | --- | --- | --- |
| **Meta-training datasets** | UK Biobank | 36,834 | 45-82 | ICA-FIX & MNI152 | 67 |
|  | ABCD | 5,985 | 9-10 | GSR & fsaverage6 | 36 |
|  | GSP | 862 | 18-35 | GSR & fsaverage6 | 23 |
|  | HBN | 930 | 5-21 | GSR & fsaverage6 | 42 |
|  | eNKI-RS | 896 | 6-85 | ICA-AROMA & MNI152 | 61 |
| **Meta-test datasets** | HCP-YA | 1,019 | 22-35 | ICA-FIX & fs_LR32k | 35 |
|  | HCP-Aging | 656 | 36-100+ | ICA-FIX & MNI152 | 45 |

We note that these datasets were opportunistically collated (e.g., by contacting potential collaborators or by downloading preprocessed data provided by the study), so the preprocessing steps varied considerably across datasets. However, we consider the heterogeneous preprocessing as a strength because the heterogeneity might help to improve (and demonstrate) generalization across preprocessing pipelines.

The phenotypes were predicted using 419 × 419 RSFC matrices, consistent with previous studies from our group (Kong et al., 2021; Chen et al., 2022; Li et al., 2022). The 419 × 419 RSFC matrices were computed using 400 cortical (Schaefer et al., 2018) and 19 subcortical parcels (Fischl et al., 2002). For each participant, RSFC was computed as the Pearson’s correlations between the average time series of each pair of brain parcels.

#### 2.1.1 UK Biobank

The UK Biobank (UKBB) dataset is a population epidemiology study with 500,000 adults (age 40-69 years) recruited between 2006 and 2010 (Sudlow et al., 2015; Miller et al., 2016). We utilized fMRI data from 36,834 participants and 67 phenotypes (selected from a total of 3,937 phenotypes) from the UK Biobank dataset. The detailed phenotypic selection procedures followed our previous study (He et al., 2022). The sample size is slightly smaller than our previous study (He et al., 2022) because of participants voluntarily withdrawing from the UK Biobank study. More specifically, ICA-FIX pre-processed volumetric rs-fMRI time series in native participant space were downloaded from the UK Biobank (Alfaro-Almagro et al., 2018). The time series were then projected to MNI152 2-mm template space, and averaged within each cortical and each subcortical parcel. Pearson’s correlations were used to generate the 419 × 419 RSFC matrices.

#### 2.1.2 ABCD

The adolescent brain cognitive development (ABCD) is a dataset of children (age 9-10 years) and a diverse set of behavioral measures (Volkow et al., 2018). We considered data from 11875 children from the ABCD 2.0.1 release. We used 36 phenotypes in total, including 16 cognitive measures, 9 personality measures, and 11 mental health measures, consistent with our previous studies (Ooi et al., 2022; Chen et al., 2023).

Details of the fMRI preprocessing can be found in previous studies (J. Chen et al., 2023; Ooi et al., 2022) but briefly, minimally preprocessed fMRI data (Hagler Jr et al., 2019) were further processed with the following steps: (1) removal of initial frames (number of frames removed depended on the type of scanner; Hagler Jr et al., 2019); (2) alignment with the T1 images using boundary-based registration (BBR; Greve & Fischl, 2009) with FsFast (http://surfer.nmr.mgh.harvard.edu/fswiki/FsFast); (3) respiratory pseudomotion motion filtering was performed by applying a bandstop filter of 0.31-0.43Hz (Fair et al., 2020) (4) functional runs with BBR costs greater than 0.6 were excluded; (5) motion correction and outlier detection: framewise displacement (FD; Jenkinson et al., 2002) and voxel-wise differentiated signal variance (DVARS; Power et al., 2012) were computed using fsl_motion_outliers. Volumes with FD > 0.3 mm or DVARS > 50, along with one volume before and two volumes after, were marked as outliers (i.e., censored frames). Uncensored segments of data containing fewer than five contiguous volumes were also censored (Gordon et al., 2016; Kong et al., 2019). BOLD runs with over half of frames censored and runs with max FD > 5mm were removed; (6) the following nuisance covariates were regressed out of the fMRI time series: a vector of ones and linear trend, global signal, six motion correction parameters, averaged ventricular signal, averaged white matter signal, and their temporal derivatives. Regression coefficients were estimated from the non-censored volumes; (7) interpolation of censored frames with Lomb-Scargle periodogram (Power et al., 2014); (8) band-pass filtering (0.009 Hz ≤ f ≤ 0.08 Hz); (9) projection onto FreeSurfer (Fischl, 2012) fsaverage6 surface space; (10) smoothing by a 6 mm full-width half-maximum (FWHM) kernel.

We also excluded participants who did not have at least 4 minutes for rs-fMRI and excluded participants without all 36 phenotypes, resulting in 5,985 participants. For each participant, the fMRI time series were averaged within each cortical parcel (in fsaverage6 surface space) and each subcortical parcel in the participant’s native volumetric space. Pearson’s correlations were used to generate the 419 × 419 RSFC matrices.

#### 2.1.3 GSP

The Brain Genomics Superstruct Project (GSP) contains fMRI and multiple behavioral measures from healthy young adults aged 18 to 35 years old (Holmes et al., 2015). We used 23 behavioral phenotypes including cognitive and personality measures, consistent with our previous study (Li et al., 2019).

Details of the fMRI preprocessing can be found in previous studies (Li et al., 2019), but briefly, the pipeline comprised the following steps: (1) removal of the first four frames; (2) slice time correction with FSL (Jenkinson et al., 2012; Smith et al., 2004) package; (3) motion correction and outlier detection: FD and DVARS were estimated using fsl_motion_outliers. Volumes with FD > 0.2mm or DVARS > 50 were marked as outliers (censored frames). One frame before and two frames after these volumes were flagged as censored frames. Uncensored segments of data lasting fewer than five contiguous volumes were also labeled as censored frames (Gordon et al., 2016). BOLD runs with more than half of the volumes labeled as censored frames were removed; (4) alignment with structural image using boundary-based registration with FsFast (Greve & Fischl, 2009); (5) regress the following nuisance regressors: a vector of ones and linear trend, six motion correction parameters, averaged white matter signal, averaged ventricular signal, mean whole brain signal, and their temporal derivatives. Regression coefficients were estimated from the non-censored volumes; (6) interpolation of censored frames with Lomb-Scargle periodogram; (7) band-pass filtering (0.009 Hz ≤ f ≤ 0.08 Hz); (8) projection onto the FreeSurfer fsaverage6 surface space; (9) smoothing with 6mm FWHM and down-sampling to fsaverage5 surface space.

We also removed participants without full 23 phenotypes, yielding 862 participants. For each participant, the fMRI time series were averaged within each cortical parcel (in fsaverage6 surface space) and each subcortical parcel in the participant’s native volumetric space. Pearson’s correlations were used to generate the 419 × 419 RSFC matrices.

#### 2.1.4 HBN

The Healthy Brain Network (HBN) contains New York area participants (age 5–21 years) with brain imaging, psychiatric, behavioral, cognitive, and lifestyle information (Alexander et al., 2017). We downloaded data from 2196 participants (HBN release 1-7). We manually selected commonly used cognitive performance scores and behavioral scores with less than 10% of missing values, resulting in 42 phenotypes.

Resting-state fMRI data were pre-processed with the following steps: (1) removal of the first 8 frames; (2) slice time correction; (3) motion correction and outlier detection: frames with FD > 0.3mm or DVARS > 60 were flagged as censored frames. 1 frame before and 2 frames after these volumes were flagged as censored frames. Uncensored segments of data lasting fewer than five contiguous frames were also labeled as censored frames. BOLD runs with over half of the frames censored and runs with max FD > 5mm were removed; (4) correcting for spatial distortion caused by susceptibility-induced off-resonance field; (5) alignment with structural image using boundary-based registration; (6) nuisance regression: regressed out a vector of ones and linear trend, global signal, six motion correction parameters, averaged ventricular signal, averaged white matter signal, and their temporal derivatives. Regression coefficients were estimated from the non-censored volumes; (7) band-pass filtering (0.009 Hz ≤ f ≤ 0.08 Hz); (8) interpolation of censored frames with Lomb-Scargle periodogram; (9) projection onto the FreeSurfer fsaverage6 surface space; (10) smoothing with 2mm FWHM and down-sampling to fsaverage5 surface space.

We excluded individuals who did not have at least 4 minutes of uncensored rs-fMRI data and removed participants with no relevant phenotypes, resulting in 930 participants. For each participant, the fMRI time series were averaged within each cortical parcel (in fsaverage6 surface space) and each subcortical parcel in the participant’s native volumetric space. Pearson’s correlations were used to generate the 419 × 419 RSFC matrices.

#### 2.1.5 eNKI-RS

The enhanced Nathan Kline Institute-Rockland Sample (eNKI-RS) is a community sample of over 1000 participants (age 6-85 years), with measures including various physiological and psychological assessments, genetic information, and neuroimaging data (Nooner et al., 2012). We manually selected commonly used cognitive performance measures and behavioral scores with less than 10% of missing value, yielding 61 phenotypes and 896 participants with at least one phenotype.

Details of the fMRI preprocessing can be found in our previous study (Wu et al., 2022), but briefly, eNKI-RS data were pre-processed with fMRIprep (Esteban et al., 2019) with default configuration and additional ICA-AROMA denoising (Pruim et al., 2015a; 2015b). Additional nuisance regression was then performed with regressors corresponding to 24 motion parameters, white matter signal, CSF signal and their temporal derivatives (Wu et al., 2022). The pre-processed fMRI data in MNI152 space were used to compute 419 × 419 RSFC matrices

#### 2.1.6 HCP-YA

The Human Connectome Project (HCP Young Adult, HCP-YA) contains brain imaging data and phenotypes from healthy young adults (age 22-35 years) (Van Essen et al., 2013). We used 35 phenotypes across cognition, personality, and emotion, consistent with our previous study (He et al., 2022). There are 1,019 participants with all 35 phenotypes in the end.

For the RSFC data, we used ICA-FIX MSMALL time series in the grayordinate (combined surface and subcortical volumetric) fsLR_32k space (Glasser et al., 2013). The time series were averaged within each cortical and each subcortical parcel to calculate 419 × 419 RSFC matrices.

#### 2.1.7 HCP-Aging

The Human Connectome Project Aging (HCP-Aging) study enrolls 1,500+ healthy adults (age 36-100+ years) (Harms et al., 2018). We manually selected commonly used behavioral measures, resulting in 45 phenotypes and 656 participants with at least one phenotype. The resting-fMRI data after ICA-FIX denoising in MNI152 space were used, following our previous study (Wu et al., 2022). Nuisance regression was then implemented, controlling for 24 motion parameters, white matter signal, CSF signal, and their temporal derivatives (Wu et al., 2022). The time series were averaged within each cortical and each subcortical parcel to calculate 419 × 419 RSFC matrices.

### 2.2 Data split overview

We split the datasets into a meta-training (source) set and a meta-test (target) set, as shown in Figure 1. For each meta-training dataset, we randomly divided the participants into training and validation sets comprising 80% and 20% of the participants respectively. The training and validation sets are used to train and tune the hyperparameters of one or more “base-learners” to predict corresponding source phenotypes from the meta-training dataset.

For each meta-test dataset, there are target phenotypes we want to predict from RSFC. For cross-dataset prediction, we trained a “meta-learner” using K participants in the meta-test dataset (i.e., K-shot, where K = 10, 20, 50, 100, 200) with observed meta-test phenotypes. The meta-learner exploits the relationship between source and target phenotypes via the previously trained base-learners from the meta-training datasets, thus transferring knowledge from the meta-training datasets to the meta-test dataset. Finally, we evaluated the prediction performance of meta-test phenotypes on the remaining N – K meta-test participants, using Pearson’s correlation and predictive coefficient of determinant (COD) as metrics.

### 2.3 Prediction approaches

Across all approaches, we vectorized the lower triangular entries of each 419 × 419 RSFC matrix into a feature vector (i.e., 87571 × 1 vector) to predict phenotypic measures. We note that certain datasets were processed with global signal regression (GSR), while others were processed with ICA-FIX (Table 1). It is well-known that GSR centers the distribution of RSFC values at zero (Murphy et al., 2009), which is not the case for ICA-FIX. Therefore, for all cross-dataset algorithms (i.e., all algorithms except kernel ridge regression), we normalized the RSFC vector for each participant independently, by subtracting the mean and then dividing by the L2-norm of the 87571 × 1 FC vector.

Following our previous study (He et al., 2022), statistical difference between algorithms was evaluated using a bootstrapping approach (more details in Supplementary Methods S3). Multiple comparisons were corrected using a false discovery rate (FDR) of q < 0.05. FDR was applied to all K-shots, across all pairs of algorithms and both evaluation metrics (Pearson’s correlation and COD).

#### 2.3.1 Baseline 1: Classical KRR

We choose kernel ridge regression (KRR; Figure 2A) as a baseline algorithm that does not utilize meta-training on the meta-training set. KRR has been shown to be a highly competitive algorithm for MRI prediction of phenotypic measures (He et al., 2020; Ooi et al., 2022; Kong et al., 2023). The procedure is as follows. Suppose the meta-test dataset has N participants in total. For each target phenotype in the meta-test dataset, we trained a KRR and tuned the hyper-parameter λ (L2 regularization weight) with 5-fold cross-validation, using K random participants with observed target phenotypes (i.e., K-shot). The optimal λ was then used to train a final KRR model using all K participants. We then evaluated the model performance on the remaining N – K participants using Pearson’s correlation and COD. The procedure was repeated 100 times with a different random set of K participants. The evaluation metrics were averaged across the 100 repetitions to ensure the robustness of the results.

**Figure 2.**
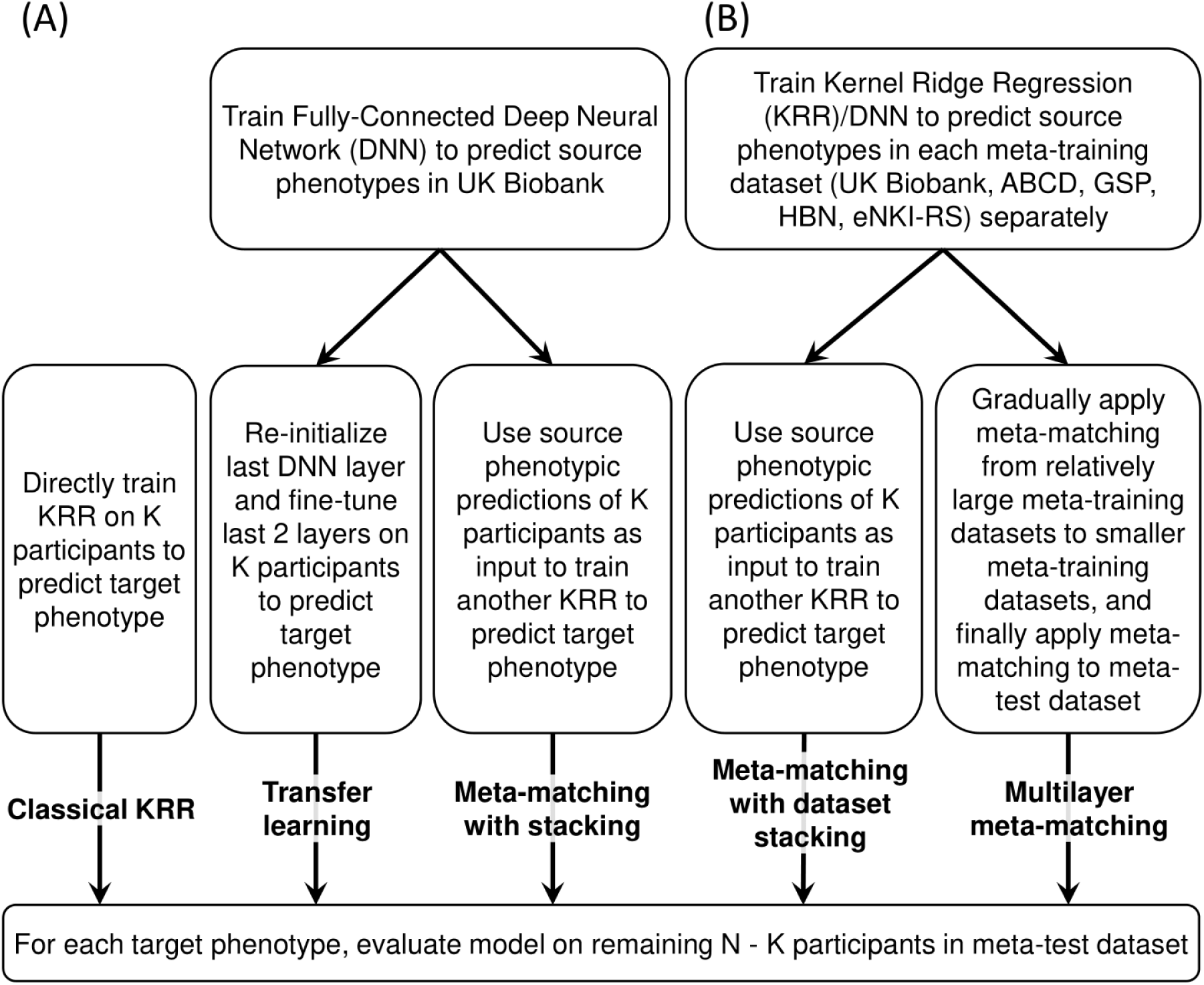
Schematic of different approaches. (A) Schematic of three baselines: classical kernel ridge regression (KRR), transfer learning, and meta-matching with stacking from our previous study (He et al., 2022). (B) Schematic of two proposed approaches: meta-matching with dataset stacking and multilayer meta-matching. Observe the large sample imbalance in the meta-training set with the smallest source dataset comprising 862 participants and the largest source dataset comprising 36,834 participants.

#### 2.3.2 Baseline 2: Transfer learning

As a second baseline, we consider transfer learning (Weiss et al., 2016). As illustrated in Figure 2A, we pre-trained a deep neural network (DNN) in the UK Biobank to simultaneously predict 67 source phenotypes from RSFC (maximum training epochs = 100). The DNN is a simple fully-connected feedforward neural network (also known as a multi-layer perceptron) with 67 output nodes. Rectifying linear units (ReLU) were used as activation functions for all hidden layers. As mentioned in Section 2.2, 80% of the data was used for training and 20% was used for tuning DNN hyper-parameters. The hyper-parameters (e.g., number of layers, number of nodes, learning rate, dropout rate, etc.) were tuned using the Optuna package (Akiba et al., 2019). Detailed information about DNN hyper-parameters is found in Supplementary Methods S1.

The pre-trained DNN was then translated using K meta-test participants to predict a target phenotype. Because we are predicting different phenotypes in the meta-test dataset, for a given target phenotype, the last layer of the pre-trained DNN was re-initialized from scratch, and the last two layers of the DNN were then fine-tuned on K random participants with observed target phenotypes (i.e., K-shot). An optimal fixed learning rate was obtained by 5-fold cross-validation and grid search of the K participants. The optimal learning rate was then used to perform fine-tune a final model using all K participants. For both the 5-fold cross validation and the final round of fine-tuning, the maximum fine-tuning epochs was set to be 10 with 80% of K participants used for training and 20% used to evaluate validation loss for early stopping, to reduce the possibility of overfitting. This final trained model was evaluated in the remaining N – K participants.

#### 2.3.3 Baseline 3: Meta-matching with stacking

The third baseline is the “meta-matching with stacking” algorithm (Figure 2A) from the original meta-matching study (He et al., 2022). The original study proposed several meta-matching algorithms. Here we used the stacking approach because it exhibited the best prediction performance in the original study.

Similar to transfer learning, the meta-matching with stacking approach utilized the same pre-trained DNN from the UK Biobank (see Section 2.3.2). To adapt the DNN to the meta-test dataset, the DNN was applied to the RSFC of the K participants, yielding 67 predictions per participant. The 67 predictions were then used as features to train a KRR model for predicting the target phenotype using the K participants (i.e., stacking; Wolpert, 1992).

The KRR model utilized the correlation kernel and the KRR hyperparameter λ was tuned using grid search and 5-fold cross-validation on the K participants. The optimal λ was then used to train a final KRR model using all K participants. The prediction performances were evaluated on the remaining N – K participants using Pearson’s correlation and COD as metrics. This procedure was repeated 100 times with a different random sample of K participants.

It is worthwhile highlighting a deviation from the original meta-matching with stacking implementation (He et al., 2022). The original implementation utilized K features for stacking when K < 67. Here, we decided to simply use all 67 features because experimentation after the publication of our previous study (not shown) suggested the constraint was unnecessary.

#### 2.3.4 Meta-matching with dataset stacking

A naive approach to extending meta-matching with stacking to multiple datasets is to train independent prediction model(s) in each meta-training (source) dataset and then “stack” the prediction models based on K participants in the meta-test dataset. We refer to this approach as meta-matching with dataset stacking (Figure 2B).

For the UK Biobank, we trained a DNN model to predict 67 phenotypes, as well as 67 KRR models to predict 67 phenotypes, to improve prediction performance via ensemble learning (Dietterich, 2000), yielding 67 × 2 = 138 predictions. We note that the DNN model is identical to that from the transfer learning baseline. The remaining four datasets (ABCD, GSP, HBN, eNKI-RS) were significantly smaller than the UK Biobank, so instead of training a DNN, we simply trained a KRR model for each meta-test dataset (including the UK Biobank) and each target phenotype. The KRR and DNN models were applied to the RSFC of the K participants (of the meta-test dataset), yielding a total of 67 × 2 + 36 + 23 + 42 + 61 = 296 phenotypic predictions for each participant.

Similar to the meta-matching with stacking approach (Section 2.3.3), the predictions were then used as features to train a KRR model for predicting the target phenotype using the K participants (i.e., stacking). The KRR model utilized the correlation kernel and the KRR hyperparameter λ was tuned using grid search and 5-fold cross-validation on the K participants. The optimal λ was then used to train a final KRR model using all K participants.

The prediction performances were evaluated on the remaining N – K participants using Pearson’s correlation and COD as metrics. This procedure was repeated 100 times with a different random sample of K participants.

#### 2.3.5 Multilayer meta-matching

As an alternative to “meta-matching with dataset stacking”, we made use of the fact “meta-matching with stacking” can improve the prediction of smaller datasets. Therefore, “multilayer meta-matching” (Figure 2B) gradually applied meta-matching with stacking from relatively large source datasets (e.g., UK Biobank) to smaller datasets (e.g., GSP, HBN, etc), to generate additional features for a final round of stacking using the K participants from the meta-test dataset.

In the current study, we instantiated multilayer meta-matching by dividing the meta-training datasets into three groups: extra-large source dataset (comprising only UK Biobank in the current study), large source datasets (comprising only ABCD in the current study) and medium size datasets (comprising GSP, HBN and eNKI-RS in the current study). Multilayer meta-matching proceeds as follows (Figure 3).

**Figure 3.**
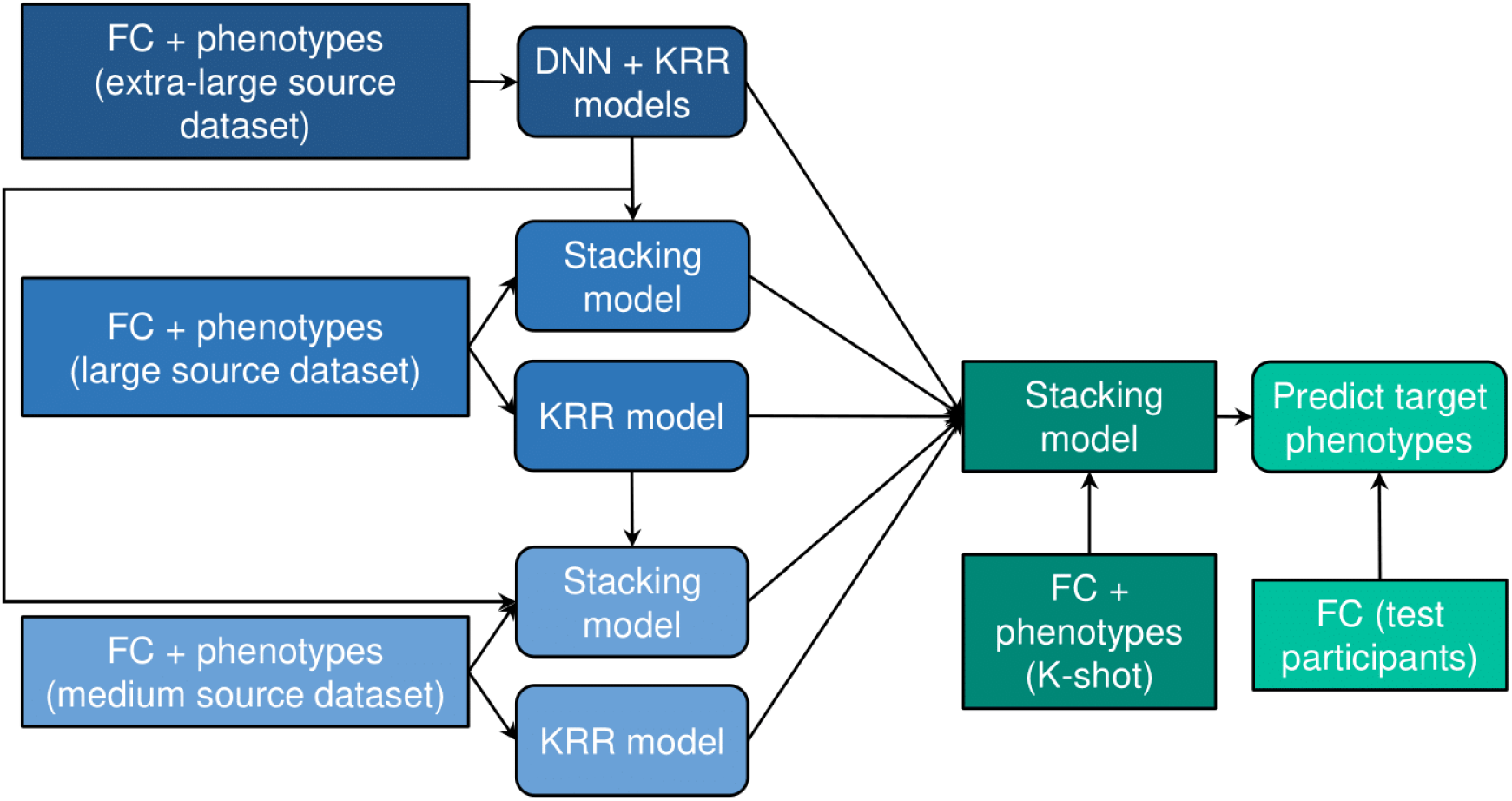
Multilayer meta-matching. We divided source datasets into extra-large (UK Biobank), large (ABCD), and medium (GSP/HBN/eNKI) source datasets. Multi-layer meta-matching gradually applied meta-matching with stacking from relatively large source datasets (e.g., UK Biobank) to smaller datasets (e.g., HCP), to generate additional features for a final round of stacking using the K participants from the meta-test dataset.

In the case of the extra-large dataset (UK Biobank), we have previously trained DNN and KRR models to predict 67 phenotypes (Section 2.3.4). The same two models were applied to the K meta-test dataset participants, yielding 67 × 2 = 134 phenotypic predictions, which will be concatenated with the predictions from the other models (below) for stacking.

In the case of the large dataset (ABCD), we have previously trained a KRR model to predict 36 phenotypes in the ABCD dataset (Section 2.3.4). The same model was applied to the K meta-test dataset participants, yielding 36 predictions. Furthermore, the DNN and KRR models from the extra-large dataset (UK Biobank) were also combined to predict the 36 ABCD phenotypes via the meta-matching with stacking procedure (He et al., 2022). The resulting stacking model was applied to the K meta-test dataset participants, yielding 36 predictions. Therefore, models from the ABCD dataset yielded a total of 36 × 2 = 72 phenotypic predictions for each of the K meta-test dataset participants, which will be concatenated with the 134 predictions from the UK Biobank (above) and predictions from the other models (below) for stacking.

Finally, in the case of the medium source dataset (GSP, HBN or eNKI-RS), let us use the GSP dataset, which had 23 phenotypes, as an example. First, we have previously trained a KRR model to predict 23 phenotypes in the GSP dataset (Section 2.3.4). The same model was applied to the K meta-test dataset participants, yielding 23 predictions. Second, the DNN and KRR models from the extra-large dataset (UK Biobank), as well as the KRR models from the large dataset (ABCD) were also combined to predict the 23 GSP phenotypes via the meta-matching with stacking procedure (He et al., 2022). The resulting stacking model was applied to the K meta-test dataset participants, yielding 23 predictions. Therefore, in total, the GSP dataset contributed 23 × 2 = 46 phenotypic predictions in each of the K meta-test dataset participants. Similarly, the HBN and eNKI-RS datasets contributed 42 × 2 = 84 and 61 × 2 = 122 phenotypic predictions.

Finally, all the phenotypic predictions (134 + 72 + 46 + 84 + 122 = 458) were concatenated and used to train a KRR model on the K meta-test dataset participants (i.e., stacking). Once again, the KRR model utilized the correlation kernel and the KRR hyperparameter λ was tuned using grid search and 5-fold cross-validation on the K participants. The optimal λ was then used to train a final KRR model using all K participants.

The prediction performances were evaluated on the remaining N – K participants using Pearson’s correlation and COD as metrics. This procedure was repeated 100 times with a different random sample of K participants.

### 2.4 Feature importance based on the Haufe transform

Although meta-matching improved phenotypic prediction performance, a question is whether the interpretation of the resulting models is biased by pre-trained prediction models. Here, we applied the Haufe transform for each approach in the K = 100 scenario, which involved computing the covariance between each FC edge and the phenotypic prediction (of the mode) across the K participants (Haufe et al., 2014; J. Chen et al., 2022). The result is a feature importance value for each RSFC edge. A positive (or negative) feature importance value indicates that higher RSFC for the edge was associated with the prediction model predicting greater (or lower) value for the phenotype. Previous studies have suggested that the Haufe transform yielded significantly more reliable feature importance values than the prediction model parameters or weights (Tian & Zalesky, 2021; Chen, Ooi et al., 2023)

Pseudo ground truth feature importance was obtained by training a KRR model on the full HCP-YA (or HCP-Aging) dataset and then applying the Haufe transform to the KRR model. In the case of classical KRR, we trained the KRR model on 100 HCP-YA (or HCP-Aging) participants and then computed the feature importance using the Haufe transform. In the case of the cross-dataset algorithms (transfer learning, meta-matching with stacking, meta-matching with dataset stacking, and multilayer meta-matching), we translated the models (trained on source datasets) on the 100 HCP-YA (or HCP-Aging) participants and then computed the feature importance.

We then correlated the resulting feature importance values with the pseudo ground truth. We repeated this procedure 100 times, and averaged the correlations with the pseudo ground truth across 100 repetitions.

### 2.5 Data and code availability

This study utilized publicly available data from the UK Biobank (https://www.ukbiobank.ac.uk/), ABCD (https://nda.nih.gov/study.html?id=824), GSP (http://neuroinformatics.harvard.edu/gsp/), HBN (https://fcon_1000.projects.nitrc.org/indi/cmi_healthy_brain_network), eNKI-RS (http://fcon_1000.projects.nitrc.org/indi/enhanced/) and HCP (https://www.humanconnectome.org/). Data can be accessed via data use agreements.

Code for the classical (KRR) baseline and meta-matching algorithms can be found here (https://github.com/ThomasYeoLab/CBIG/tree/master/stable_projects/predict_phenotypes/Ch <u>en2024_MMM</u>). The trained models for multilayer meta-matching are also publicly available (GITHUB_LINK). The code was reviewed by one of the co-authors (LA) before merging into the GitHub repository to reduce the chance of coding errors.

## 3. Results

### 3.1 Meta-matching with stacking outperformed classical KRR and transfer learning

Figures 4A and 4B show the prediction accuracy (Pearson’s correlation coefficient) of various approaches in the HCP-YA and HCP-Aging meta-test datasets respectively. Results were averaged across 35 HCP-YA (or 45 HCP-Aging) phenotypes. The horizontal axis is the number of few-shot participants (K, where K = 10, 20, 50, 100, 200). The vertical axis is Pearson’s correlation of phenotypic prediction. Boxplots represent variability across the 100 repetitions of sampling K participants (i.e., K-shot). Figure 5 shows results for COD. Bootstrapping results are shown in Figures S1 and S2, while p values are reported in Tables 2 and 3. All bolded p values (Tables 2 and 3) survived an FDR of q < 0.05.

Consistent with our previous study (He et al., 2022), meta-matching with stacking outperformed classical KRR in the HCP-YA dataset (Figures 4A and 5A; Tables 2). Here, we extended the previous results by showing consistent improvements over KRR in the HCP-Aging dataset.

**Figure 4.**
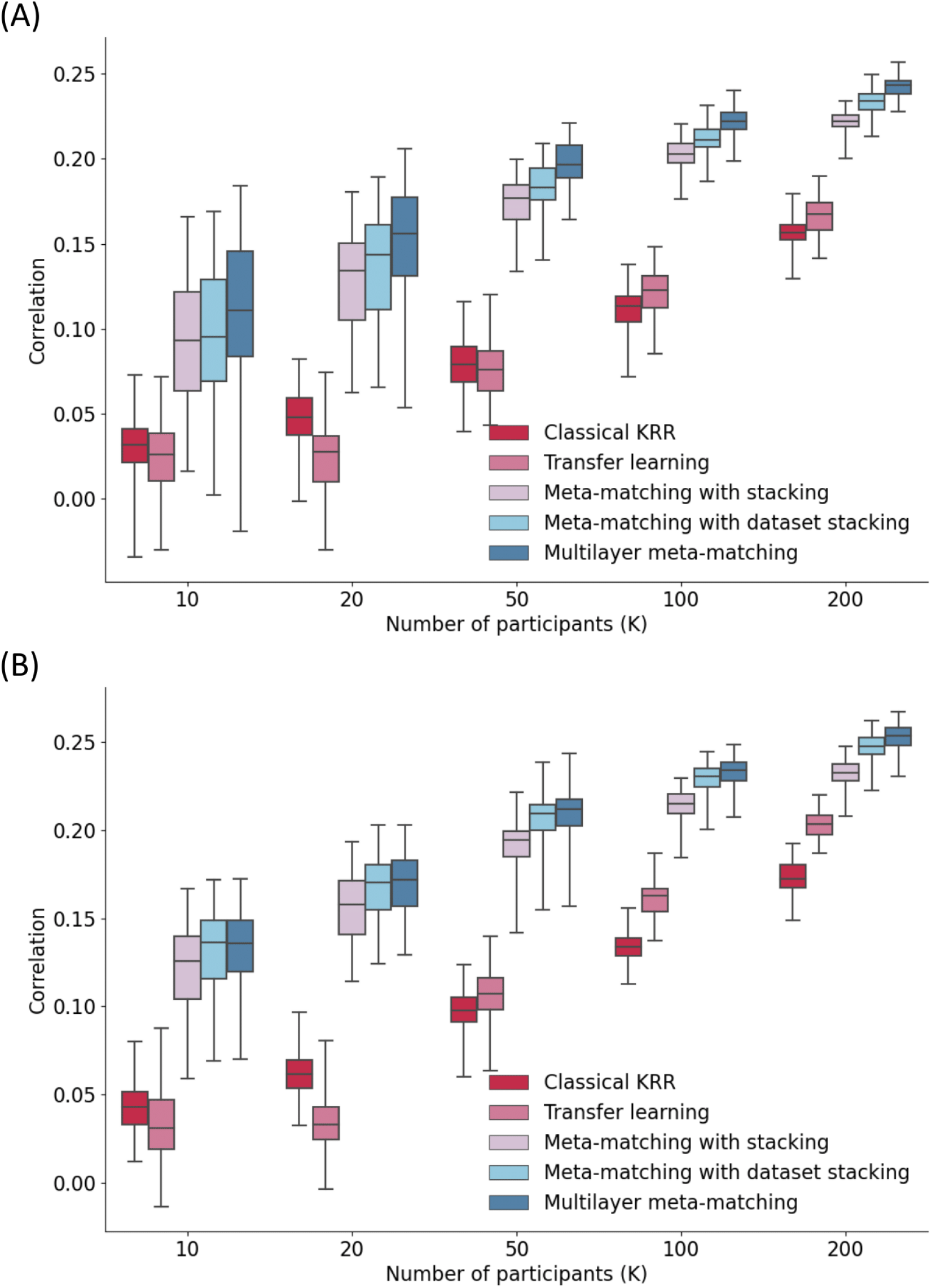
Prediction performance (Pearson’s correlation) in the HCP-YA and HCP-Aging datasets. (A) Phenotypic prediction performance in terms of Pearson’s correlation (averaged across 35 meta-test phenotypes) in the HCP-YA dataset. Horizontal axis is the number of participants in the HCP-YA dataset used to adapt the models trained from the meta-training source datasets. Boxplots represent variability across 100 repetitions of sampling K participants. The bottom and top edges of the box indicate the 25th and 75th percentiles, respectively. Whiskers correspond to 1.5 times the interquartile range. (B) Same plot as panel A except that the analyses were performed in the HCP-Aging dataset. Boxplots represent variability across 100 repetitions of sampling K participants. The bottom and top edges of the box indicate the 25th and 75th percentiles, respectively. Whiskers correspond to 1.5 times the interquartile range. (B) Same plot as panel A, except that the analyses were performed in the HCP-Aging dataset.

**Figure 5.**
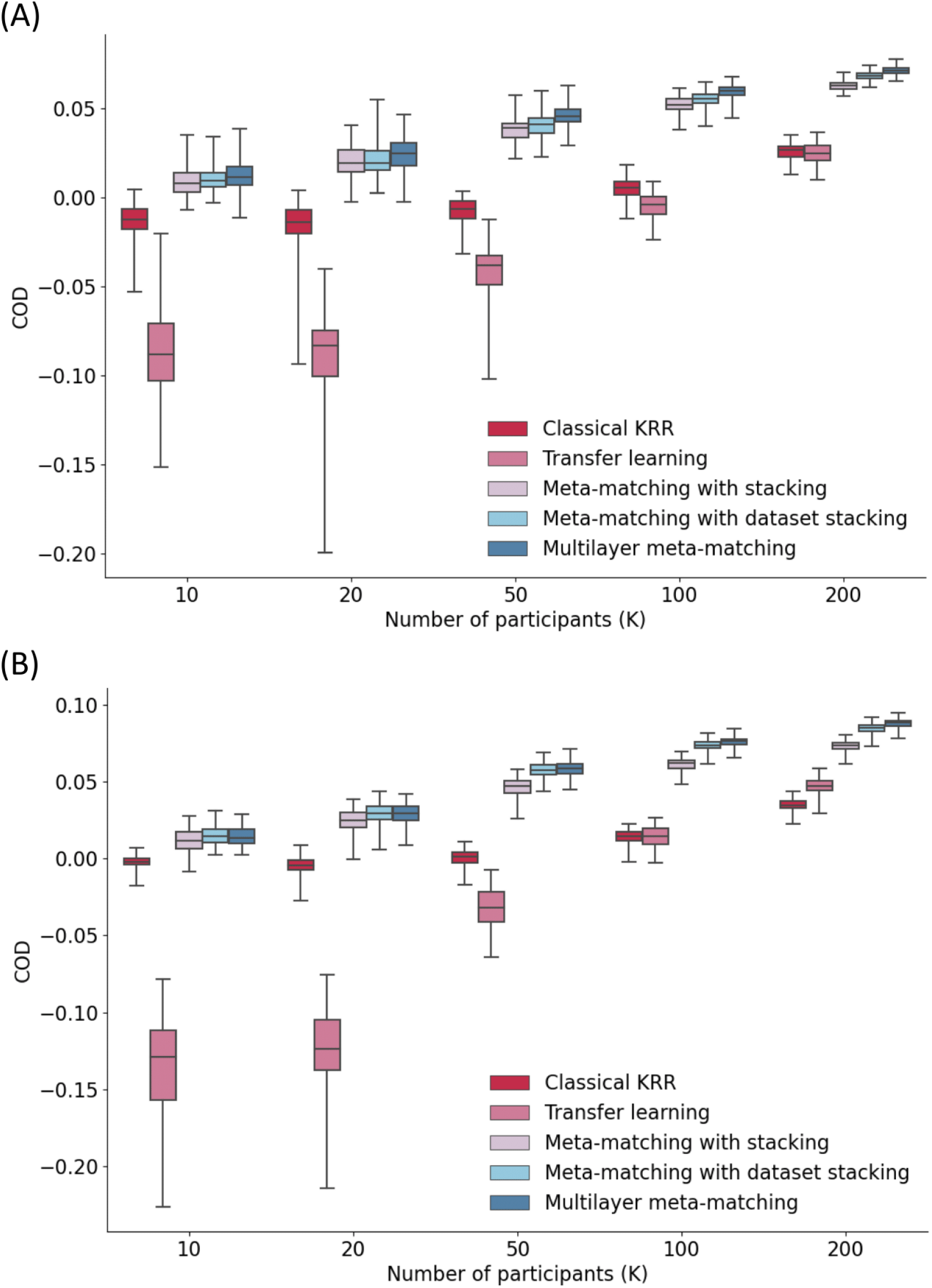
Prediction performance (COD) in the HCP-YA and HCP-Aging datasets. (A) Phenotypic prediction performance in terms of COD (averaged across 35 meta-test phenotypes) in the HCP-YA meta-test set. Horizontal axis is the number of participants in the HCP-YA dataset used to adapt the models trained from the meta-training source datasets.

More specifically, in the case of the HCP-YA dataset and K > 10 (Table 2), meta-matching with stacking was statistically better than classical KRR with largest p < 0.005 across both evaluation metrics (Pearson’s correlation and COD). In the case of HCP-Aging and K > 10 (Table 3), meta-matching with stacking was statistically better than classical KRR with largest p < 0.001 across both evaluation metrics.

**Table 2.** Statistical differences in prediction accuracy in terms of Pearson’s correlation (upper) and COD (bottom) between all pairs of approaches in the HCP-YA meta-test dataset. Here ‘MM’ stands for ‘meta-matching’ and ‘w/’ is short for ‘with’. Each cell contains five p values, corresponding to K = 10, 20, 50, 100 and 200 respectively. Bolded p values are statistically significant after FDR correction with q < 0.05.

|  |  |  |  |  |  |  |
| --- | --- | --- | --- | --- | --- | --- |
| Correlation |  | Classical KRR | Transfer learning | MM w/ stacking | MM w/ dataset stacking | Multilayer MM |
|  | Classical KRR | - | 0.787<br>0.406<br>0.898<br>0.660<br>0.295 | **0.0270**<br>**0.00247**<br>**1.13e-6**<br>**1.80e-8**<br>**4.89e-8** | **0.0134**<br>**2.64e-4**<br>**2.72e-10**<br>**3.77e-14**<br>**1.11e-16** | **0.00813**<br>**2.96e-5**<br>**6.86e-13**<br>**1.22e-15**<br>**0** |
|  | Transfer learning | 0.787<br>0.406<br>0.898<br>0.660<br>0.295 | - | **0.0248**<br>**0.00137**<br>**1.48e-6**<br>**9.92e-8**<br>**2.77e-6** | **0.0132**<br>**1.75e-4**<br>**3.00e-9**<br>**4.76e-12**<br>**1.01e-13** | **0.00911**<br>**3.30e-5**<br>**3.86e-11**<br>**3.44e-13**<br>**4.00e-15** |
|  | MM w/ stacking | **0.0270**<br>**0.00247**<br>**1.13e-6**<br>**1.80e-8**<br>**4.89e-8** | **0.0248**<br>**0.00137**<br>**1.48e-6**<br>**9.92e-8**<br>**2.77e-6** | - | 0.407<br>0.221<br>**0.0319**<br>**8.78e-4**<br>**2.96e-6** | 0.190<br>0.0411<br>**0.00203**<br>**6.00e-5**<br>**1.29e-7** |
|  | MM w/ dataset stacking | **0.0134**<br>**2.64e-4**<br>**2.72e-10**<br>**3.77e-14**<br>**1.11e-16** | **0.0132**<br>**1.75e-4**<br>**3.00e-9**<br>**4.76e-12**<br>**1.01e-13** | 0.407<br>0.221<br>**0.0319**<br>**8.78e-4**<br>**2.96e-6** | - | 0.182<br>0.0395<br>**0.00990**<br>**0.00867**<br>**0.00823** |
|  | Multilayer MM | **0.00813**<br>**2.96e-5**<br>**6.86e-13**<br>**1.22e-15**<br>**0** | **0.00911**<br>**3.30e-5**<br>**3.86e-11**<br>**3.44e-13**<br>**4.00e-15** | 0.190<br>0.0411<br>**0.00203**<br>**6.00e-5**<br>**1.29e-7** | 0.182<br>0.0395<br>**0.00990**<br>**0.00867**<br>**0.00823** | - |
| COD |  | Classical KRR | Transfer learning | MM w/ stacking | MM w/ dataset stacking | Multilayer MM |
|  | Classical KRR | - | **0.00438**<br>**0.00290**<br>**0.127**<br>**0.945**<br>**0.0384** | 0.0836<br>**0.00902**<br>**1.46e-4**<br>**3.56e-6**<br>**4.20e-6** | 0.0466<br>**0.00482**<br>**2.93e-5**<br>**4.84e-8**<br>**5.55e-8** | **0.0359**<br>**0.00285**<br>**4.20e-6**<br>**5.70e-9**<br>**2.32e-8** |
|  | Transfer learning | **0.00438**<br>**0.00290**<br>**0.127**<br>**0.945**<br>**0.0384** | - | **1.17e-4**<br>**1.30e-6**<br>**2.34e-8**<br>**2.03e-10**<br>**3.43e-10** | **8.53e-5**<br>**6.50e-7**<br>**9.46e-10**<br>**5.65e-14**<br>**0** | **5.99e-5**<br>**3.93e-7**<br>**1.38e-10**<br>**4.55e-15**<br>**0** |
|  | MM w/ stacking | 0.0836<br>**0.00902**<br>**1.46e-4**<br>**3.56e-6**<br>**4.20e-6** | **1.17e-4**<br>**1.30e-6**<br>**2.34e-8**<br>**2.03e-10**<br>**3.43e-10** | - | 0.676<br>0.499<br>0.104<br>**0.00163**<br>**4.29e-7** | 0.396<br>0.173<br>**0.00646**<br>**4.823e-5**<br>**4.734e-8** |
|  | MM w/ dataset stacking | 0.0466<br>**0.00482**<br>**2.93e-5**<br>**4.84e-8**<br>**5.55e-8** | **8.53e-5**<br>**6.50e-7**<br>**9.46e-10**<br>**5.65e-14**<br>**0** | 0.676<br>0.499<br>0.104<br>**0.00163**<br>**4.29e-7** | - | 0.357<br>0.127<br>**0.00858**<br>**0.0146**<br>0.0492 |
|  | Multilayer MM | **0.0359**<br>**0.00285**<br>**4.20e-6**<br>**5.70e-9**<br>**2.32e-8** | **5.99e-5**<br>**3.93e-7**<br>**1.38e-10**<br>**4.55e-15**<br>**0** | 0.396<br>0.173<br>**0.00646**<br>**4.823e-5**<br>**4.734e-8** | 0.357<br>0.127<br>**0.00858**<br>**0.0146**<br>0.0492 | - |

**Table 3.** Statistical differences in prediction accuracy in terms of Pearson’s correlation (upper) and COD (bottom) between all pairs of approaches in the HCP-Aging meta-test dataset. Here ‘MM’ stands for ‘meta-matching’, and ‘w/’ is short for ‘with’. Each cell contains five p values, corresponding to K = 10, 20, 50, 100 and 200 respectively. Bolded p values are statistically significant after FDR correction with q < 0.05.

|  |  |  |  |  |  |  |
| --- | --- | --- | --- | --- | --- | --- |
| Correlation |  | Classical KRR | Transfer learning | MM w/ stacking | MM w/ dataset stacking | Multilayer MM |
|  | Classical KRR | - | 0.580<br>0.227<br>0.652<br>**0.0215**<br>**1.86e-4** | **5.61e-4**<br>**2.30e-6**<br>**7.59e-9**<br>**2.97e-10**<br>**2.88e-9** | **4.42e-5**<br>**9.44e-9**<br>**3.02e-13**<br>**0**<br>**0** | **1.48e-5**<br>**9.80e-10**<br>**1.64e-14**<br>**0**<br>**0** |
|  | Transfer learning | 0.580<br>0.227<br>0.652<br>**0.0215**<br>**1.86e-4** | - | **0.00109**<br>**1.67e-6**<br>**4.11e-7**<br>**1.04e-5**<br>**0.00972** | **2.42e-4**<br>**4.38e-8**<br>**3.70e-10**<br>**1.94e-10**<br>**1.68e-7** | **1.40e-4**<br>**1.41e-8**<br>**5.85e-11**<br>**3.17e-11**<br>**1.46e-8** |
|  | MM w/ stacking | **5.61e-4**<br>**2.30e-6**<br>**7.59e-9**<br>**2.97e-10**<br>**2.88e-9** | **0.00109**<br>**1.67e-6**<br>**4.11e-7**<br>**1.04e-5**<br>**0.00972** | - | 0.278<br>0.0938<br>**0.00280**<br>**3.59e-5**<br>**1.91e-6** | 0.233<br>0.0715<br>**0.00196**<br>**1.90e-5**<br>**1.05e-7** |
|  | MM w/ dataset stacking | **4.42e-5**<br>**9.44e-9**<br>**3.02e-13**<br>**0**<br>**0** | **2.42e-4**<br>**4.38e-8**<br>**3.70e-10**<br>**1.94e-10**<br>**1.68e-7** | 0.278<br>0.0938<br>**0.00280**<br>**3.59e-5**<br>**1.91e-6** | - | 0.463<br>0.321<br>0.182<br>0.0826<br>**0.00728** |
|  | Multilayer MM | **1.48e-5**<br>**9.80e-10**<br>**1.64e-14**<br>**0**<br>**0** | **1.40e-4**<br>**1.41e-8**<br>**5.85e-11**<br>**3.17e-11**<br>**1.46e-8** | 0.233<br>0.0715<br>**0.00196**<br>**1.90e-5**<br>**1.05e-7** | 0.463<br>0.321<br>0.182<br>0.0826<br>**0.00728** | - |
| COD |  | Classical KRR | Transfer learning | MM w/ stacking | MM w/ dataset stacking | Multilayer MM |
|  | Classical KRR | - | **7.61e-5**<br>**2.27e-5**<br>**0.0573**<br>**0.287**<br>**3.79e-7** | 0.0807<br>**9.83e-4**<br>**8.85e-9**<br>**9.85e-11**<br>**2.51e-14** | **0.0230**<br>**4.71e-5**<br>**1.34e-13**<br>**0**<br>**0** | **0.0215**<br>**3.58e-5**<br>**3.04e-14**<br>**0**<br>**0** |
|  | Transfer learning | **7.61e-5**<br>**2.27e-5**<br>**0.0573**<br>**0.287**<br>**3.79e-7** | - | **8.36e-6**<br>**6.08e-8**<br>**1.05e-7**<br>**4.24e-8**<br>**2.44e-4** | **7.07e-6**<br>**3.12e-8**<br>**1.69e-9**<br>**3.19e-13**<br>**8.85e-11** | **7.35e-6**<br>**3.07e-8**<br>**9.89e-10**<br>**6.42e-14**<br>**6.49e-12** |
|  | MM w/ stacking | 0.0807<br>**9.83e-4**<br>**8.85e-9**<br>**9.85e-11**<br>**2.51e-14** | **8.36e-6**<br>**6.08e-8**<br>**1.05e-7**<br>**4.24e-8**<br>**2.44e-4** | - | 0.611<br>0.241<br>**6.64e-4**<br>**4.07e-7**<br>**2.83e-9** | 0.655<br>0.250<br>**4.55e-4**<br>**5.24e-8**<br>**4.44e-11** |
|  | MM w/ dataset stacking | **0.0230**<br>**4.71e-5**<br>**1.34e-13**<br>**0**<br>**0** | **7.07e-6**<br>**3.12e-8**<br>**1.69e-9**<br>**3.19e-13**<br>**8.85e-11** | 0.611<br>0.241<br>**6.64e-4**<br>**4.07e-7**<br>**2.83e-9** | - | 0.987<br>0.791<br>0.229<br>**0.0312**<br>**9.76e-4** |
|  | Multilayer MM | **0.0215**<br>**3.58e-5**<br>**3.04e-14**<br>**0**<br>**0** | **7.35e-6**<br>**3.07e-8**<br>**9.89e-10**<br>**6.42e-14**<br>**6.49e-12** | 0.655<br>0.250<br>**4.55e-4**<br>**5.24e-8**<br>**4.44e-11** | 0.987<br>0.791<br>0.229<br>**0.0312**<br>**9.76e-4** | - |

Furthermore, meta-matching with stacking also outperformed transfer learning across both datasets (Figures 4A and 5A). In the case of the HCP-YA dataset and K **≥** 10 (Table 2), meta-matching with stacking was statistically better than transfer learning with p values < 0.025 across both evaluation metrics (Pearson’s correlation and COD). In the case of HCP-Aging and K **≥** 10 (Table 3), meta-matching with stacking was statistically better than transfer learning with largest p < 0.002 across both evaluation metrics.

Interestingly, transfer learning performed consistently worse than classical KRR for K < 50, especially for the COD metric (Figures 4A and 5A; Tables 2 and 3).

### 3.2 Improvement from additional meta-training source datasets

By including additional meta-training datasets, meta-matching with dataset stacking and multilayer meta-matching were numerically better than meta-matching with stacking (which only utilized the UK Biobank) for almost all values of K (Figures 4 and 5).

In the case of the HCP-YA dataset and K > 20 (Table 2), meta-matching with dataset stacking was statistically better than meta-matching with stacking with largest p < 0.03 across both evaluation metrics (Pearson’s correlation and COD). In the case of the HCP-Aging and K > 20 (Table 3), meta-matching with dataset stacking was statistically better than meta-matching with stacking with largest p < 0.003 across both evaluation metrics.

On the other hand, in the case of the HCP-YA dataset and K > 20 (Table 2), multilayer meta-matching was statistically better than meta-matching with stacking with largest p < 0.01 across both evaluation metrics. In the case of the HCP-Aging and K > 20 (Table 3), multilayer meta-matching was statistically better than meta-matching with stacking with largest p < 0.002 across both evaluation metrics.

We observe that the p values for multilayer meta-matching were generally stronger (i.e., smaller) than meta-matching with dataset stacking and will directly compare the two meta-matching variants in the next section.

### 3.3 Multilayer meta-matching modestly outperformed meta-matching with dataset stacking

Multi-layer meta-matching was numerically better than meta-matching with dataset stacking for almost all values of K. This improvement was significant for larger values of K.

In the case of the HCP-YA dataset and K > 20 (Table 2), multi-layer meta-matching was statistically better than meta-matching with dataset stacking with largest p < 0.01 for both evaluation metrics (correlation and COD). For HCP-Aging, multilayer meta-matching was statistically better than meta-matching with dataset stacking for K = 200 for both evaluation metrics (p < 0.01; Table 3).

Overall, the results suggest that multilayer meta-matching was modestly more effective than meta-matching with dataset stacking at handling sample size imbalance among meta-training source datasets.

### 3.4 Different improvements on different phenotypes by multilayer meta-matching

Figure 6 illustrates the 100-shot prediction performance (Pearson’s correlation coefficient) of three example meta-test phenotypes across all approaches in the HCP-YA (Figure 6A) and HCP-Aging (Figure 6B) datasets. For three illustrated HCP-YA phenotypes (“Delay Discounting”, “Manual Dexterity”, “Arithmetic”), multilayer meta-matching exhibited numerically the best results. On the other hand, among the three illustrated HCP-Aging phenotypes, multilayer meta-matching was numerically worse than meta-matching with stacking and meta-matching with dataset stacking in the case of “Walking Endurance”, but was numerically the best for “MOCA score” and “Perceived Hostility”.

**Figure 6.**
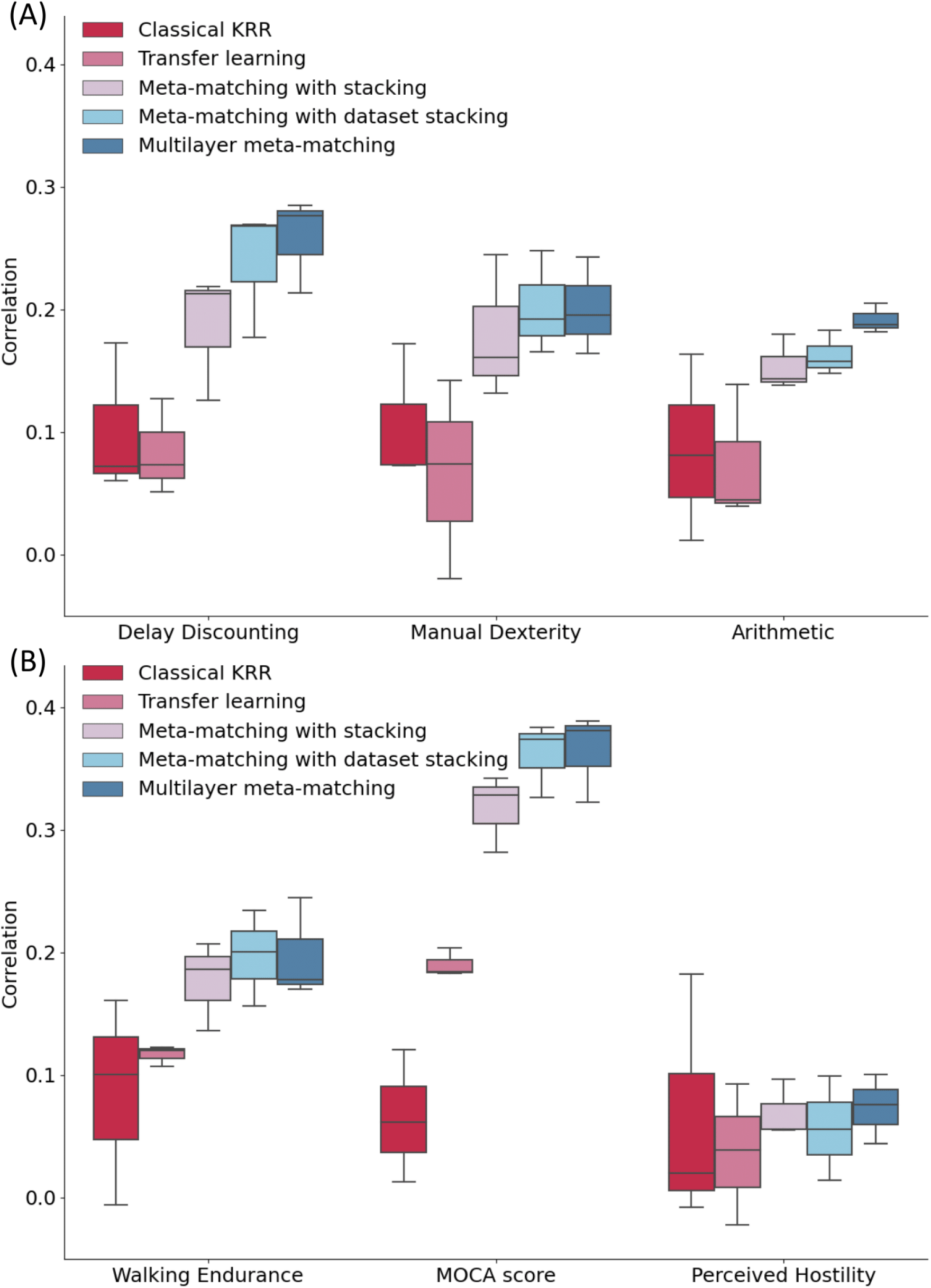
Examples of phenotypic prediction performance in the (A) HCP-YA and (B) HCP-Aging datasets in the case of 100-shot learning (K = 100). Here, prediction performance was measured using Pearson’s correlation. For each box plot, the horizontal line indicates the median. The bottom and top edges of the box indicate the 25th and 75th percentiles, respectively. Whiskers correspond to 1.5 times the interquartile range.

### 3.5 Feature importance using the Haufe transform

As shown in Figure 7, across both HCP-YA and HCP-Aging datasets, feature importance values of all three meta-matching approaches and classical KRR were equally similar to the pseudo ground truth feature importance values. On the other hand, feature importance values from transfer learning were the most different from the pseudo ground truth.

**Figure 7.**
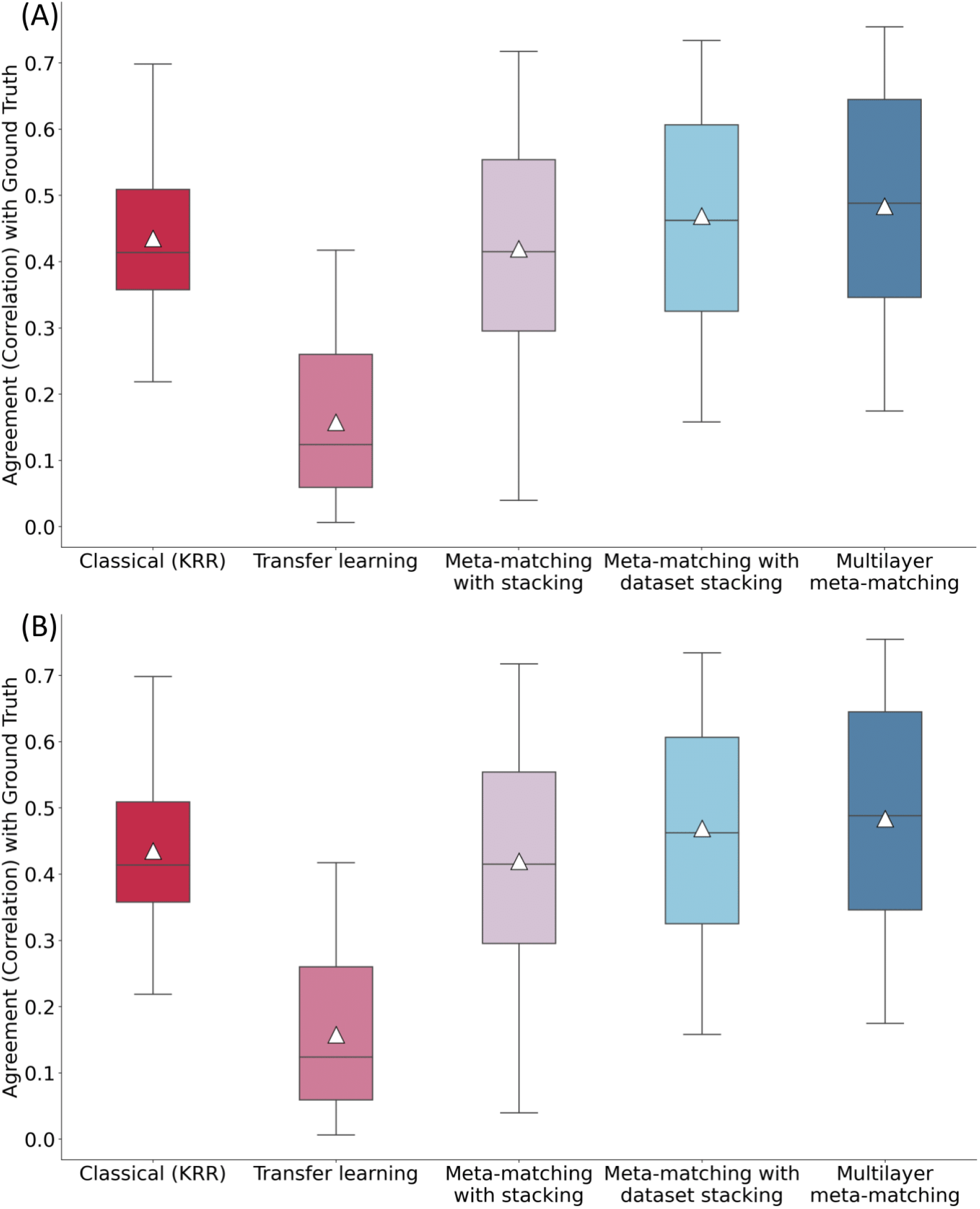
Agreement (correlation) of feature importance values with pseudo ground truth in the (A) HCP-YA and (B) HCP-Aging datasets. For each approach, the Haufe transform was used to estimate feature importance in the 100-shot scenario (K = 100), which was then compared with the pseudo ground truth. Pseudo ground truth feature importance was generated by applying the Haufe transform to a KRR model trained from the full target dataset. For each box plot, the horizontal line indicates the median, and the triangle indicates the mean. The bottom and top edges of the box indicate the 25th and 75th percentiles, respectively. Whiskers correspond to 1.5 times the interquartile range.

## 4. Discussion

In this study, we proposed two meta-matching algorithms to translate phenotypic prediction models from source datasets with disparate sizes to predict new phenotypes in small datasets. Both approaches outperformed meta-matching using a single source dataset (UK Biobank). Both approaches also outperformed classical KRR and classical transfer learning by a big margin. Furthermore, multilayer meta-matching compared favorably with meta-matching with dataset stacking across both HCP-YA and HCP-Aging datasets. In terms of feature importance based on the Haufe transform, we found that feature importance values of meta-matching approaches and classical KRR to be equally similar to the pseudo ground truth, while feature importance values of transfer learning were the furthest away from the pseudo ground truth.

The poorer performance of classical transfer learning was somewhat surprising but probably indicated the difficulty of finetuning so many parameters in the very small sample regime. More specifically, we note that classical transfer learning was even worse than KRR when the number of participants was less than 50. However, transfer learning started to catch up with KRR in both datasets when the number of participants was 200.

We note that in our previous study (He et al., 2022), one of the meta-matching variants “meta-matching finetune” outperformed KRR by a big margin but was slightly worse than meta-matching with stacking. Meta-matching finetune is similar to classical transfer learning in the sense that the last two layers of the DNN were finetuned. However, while transfer learning initialized the last layer of the DNN from scratch (Section 2.3.2), meta-matching finetune retained the weights leading to the output node that predicted the K meta-test participants the best (for each meta-test phenotype). This further supported the importance of the meta-matching approach.

One important limitation of meta-matching is that the magnitude of prediction improvement heavily depends on the correlations between meta-training and meta-test phenotypes (He et al., 2022). Consequently, we do not expect all meta-test phenotypes to benefit from meta-matching (Figure 6). However, it is important to note that this limitation exists for all meta-learning and transfer learning algorithms. Model transfer is easier if the source and target domains are more similar. Performance will degrade if the source and target domains are very different. This observation motivates the addition of more source datasets.

However, we note that the use of five source datasets (multi-layer meta-matching and meta-matching with dataset stacking) only modestly improved over the use of one source dataset (UK Biobank). One potential reason is that the UK Biobank was still more than four times larger than the combined sample size of the remaining four source datasets. Therefore, algorithmic innovation alone might not be sufficient to alleviate this issue.

Finally, we note that there are multiple possible extensions to the current work. For example, meta-matching can be applied to other imaging modalities, such as anatomical T1 images and diffusion MRI. The datasets in the current study comprised relatively healthy participants. Meta-matching might be potentially useful for psychiatric populations (Chopra et al., 2022). Including psychiatric datasets to the base model training might further improve generalization to new datasets by increasing the diversity of the source datasets.

## Acknowledgements

This work was supported by by the NUS Yong Loo Lin School of Medicine (NUHSRO/2020/124/TMR/LOA), the Singapore National Medical Research Council (NMRC) LCG (OFLCG19May-0035), NMRC CTG-IIT (CTGIIT23jan-0001), NMRC STaR (STaR20nov-0003), Singapore Ministry of Health (MOH) Centre Grant (CG21APR1009), the Temasek Foundation (TF2223-IMH-01), and the United States National Institutes of Health (R01MH120080 & R01MH133334). Our computational work was partially performed on resources of the National Supercomputing Centre, Singapore (https://www.nscc.sg). Any opinions, findings and conclusions or recommendations expressed in this material are those of the author(s) and do not reflect the views of the Singapore NRF, NMRC, MOH or the Temasek Foundation. Data used in this research were provided by: (1) the UK Biobank resource under application 25163; (2) the Adolescent Brain Cognitive Development^SM^ (ABCD) Study (https://abcdstudy.org), held in the NIMH Data Archive (NDA). This is a multisite, longitudinal study designed to recruit more than 10,000 children age 9-10 and follow them over 10 years into early adulthood. The ABCD Study® is supported by the National Institutes of Health and additional federal partners under award numbers U01DA041048, U01DA050989, U01DA051016, U01DA041022, U01DA051018, U01DA051037, U01DA050987, U01DA041174, U01DA041106, U01DA041117, U01DA041028, U01DA041134, U01DA050988, U01DA051039, U01DA041156, U01DA041025, U01DA041120, U01DA051038, U01DA041148, U01DA041093,

U01DA041089, U24DA041123, U24DA041147. A full list of supporters is available at https://abcdstudy.org/federal-partners.html. A listing of participating sites and a complete listing of the study investigators can be found at https://abcdstudy.org/consortium_members/. ABCD consortium investigators designed and implemented the study and/or provided data but did not necessarily participate in the analysis or writing of this report. This manuscript reflects the views of the authors and may not reflect the opinions or views of the NIH or ABCD consortium investigators. The ABCD data repository grows and changes over time.

The ABCD data used in this report came from http://dx.doi.org/10.15154/1504041; (3) the Brain Genomics Superstruct Project of Harvard University and the Massachusetts General Hospital (Principal Investigators: Randy Buckner, Joshua Roffman, and Jordan Smoller), with support from the Center for Brain Science Neuroinformatics Research Group, the Athinoula A. Martinos Center for Biomedical Imaging, and the Center for Human Genetic Research. Twenty individual investigators at Harvard and MGH generously contributed data to the overall project; (4) The HBN (http://www.healthybrainnetwork.org) and its initiatives are supported by philanthropic contributions from the following individuals, foundations and organizations: Margaret Bilotti; Brooklyn Nets; Agapi and Bruce Burkard; James Chang; Phyllis Green and Randolph Cōwen; Grieve Family Fund; Susan Miller and Byron Grote; Sarah and Geoff Gund; George Hall; Jonathan M. Harris Family Foundation; Joseph P. Healey; The Hearst Foundations; Eve and Ross Jaffe; Howard & Irene Levine Family Foundation; Rachael and Marshall Levine; George and Nitzia Logothetis; Christine and Richard Mack; Julie Minskoff; Valerie Mnuchin; Morgan Stanley Foundation; Amy and John Phelan; Roberts Family Foundation; Jim and Linda Robinson Foundation, Inc.; The Schaps Family; Zibby Schwarzman; Abigail Pogrebin and David Shapiro; Stavros Niarchos Foundation; Preethi Krishna and Ram Sundaram; Amy and John Weinberg; Donors to the 2013 Child Advocacy Award Dinner Auction; Donors to the 2012 Brant Art Auction; (5) the enhanced Nathan Kline InstituteRockland Sample (eNKI-RS) database (https://fcon_1000.projects.nitrc.org/indi/ enhanced/access.html); (6) the Human Connectome Project, the WU-Minn Consortium (principal investigators: David Van Essen and Kamil Ugurbil; 1U54MH091657) funded by the 16 NIH institutes and centers that support the NIH Blueprint for Neuroscience Research and by the McDonnell Center for Systems Neuroscience at Washington University; (7) the National Institute On Aging of the National Institutes of Health under Award Number U01AG052564. The content is solely the responsibility of the authors and does not necessarily represent the official views of the Ntional Institutes of Health. The associated study ID is 1376 (http://dx.doi.org/10.15154/1524254).

## Author Contribution

P.C., L.A., N.W., C.Z., S.Z., L.Q.R.O., R.K., J.C., J.W., S.C., D.B., S.B.E., A.J.H. and B.T.T.Y. designed the research. P.C. conducted the research. P.C., L.A., N.W., C.Z., S.Z., L.Q.R.O., R.K., J.C., J.W., S.C., D.B., S.B.E., A.J.H. and B.T.T.Y interpreted the results. P.C. and B.T.T.Y. wrote the manuscript and made the figures. P.C., L.A., C.Z. and N.W. reviewed and published the code. All authors contributed to project direction via discussion. All authors edited the manuscript.

