## Supplementary Materials for "Multilayer meta-matching: translating phenotypic prediction models from multiple datasets to small data"

This supplementary material is divided into Supplementary Methods and Supplementary Results to complement the Methods and Results sections in the main text, respectively.

### Supplementary Methods

Section S1 provides details about DNN used for transfer learning and meta-matching methods. Section S2 provides information about phenotypes used in each dataset.

*S1. Details of DNN trained on UK Biobank*

In this section, we provide further details about the DNN trained on the UK Biobank, which was used in transfer learning, meta-matching with stacking, meta-matching with dataset stacking and multilayer meta-matching. We used mean square error (MSE) as the loss function. The output layer of DNN has 67 nodes (which is the number of UK Biobank phenotypes). The maximum training epoch was set to 100. The learning rate decreases by half if validation loss has not decreased for 25 epochs. The epoch with the best COD on the validation set was chosen as the optimal epoch. Table S1 shows the final hyperparameters values of the DNN trained on UK Biobank datasets, which were automatically tuned using Optuna (Akiba et al., 2019) with 200 trials.

| **Hyper-parameters** | **Values** |
|---|---|
| Number of layers | 4 |
| Number of nodes in hidden layers | 512 – 256 – 128 |
| Dropout rate | 0.4 |
| Starting learning rate | 0.036612177992895435 |
| Weight decay rate | 7.321529434575473e-06 |
| Batch size | 128 |

**Table S1.** Final DNN hyper-parameters estimated by the Optuna package.

*S2. Details of phenotypes selected from each dataset*

The phenotypes selected for use in UK Biobank, ABCD, GSP, HBN, eNKI-RS, HCP-YA, and HCP-Aging datasets are shown in Table S2 – Table S8 respectively.

| Label | Description |
|---|---|
| #household | number of people in household |
| #Mem C1 | numeric memory principal component 1 |
| Age | age |
| Age edu | age completed full time education |
| Alcohol 1 | average monthly spirits intake |
| Alcohol 2 | average weekly champagne plus white wine intake |
| Alcohol 3 | average weekly beer plus cider intake |
| Blood C2 | blood assays principal component 2 |
| Blood C3 | blood assays principal component 3 |
| Blood C4 | blood assays principal component 4 |
| Blood C5 | blood assays principal component 5 |
| Body C1 | anthropometry principal component 1 |
| Body C2 | anthropometry principal component 2 |
| Body C3 | anthropometry principal component 3 |
| Bone C1 | bone-densitometry of heel principal component 1 |
| Bone C3 | bone-densitometry of heel principal component 3 |
| BP eye C2 | blood pressure & eye measures component 2 |
| BP eye C3 | blood pressure & eye measures principal component 3 |
| BP eye C4 | blood pressure & eye measures principal component 4 |
| BP eye C5 | blood pressure & eye measures principal component 5 |
| BP eye C6 | blood pressure & eye measures principal component 6 |
| Breath C1 | spirometry principal component 1 |
| Cancer C1 | cancer principal component 1 |
| Carotid C1 | carotid ultrasound principal component 1 |
| Carotid C5 | carotid ultrasound principal component 5 |
| Deprive C1 | multiple deprivation principal component 1 |
| Digit 1 | symbol digit substitution principal component 1 |
| Digit-o C1 | symbol digit substitution online principal component 1 |
| Digit-o C6 | symbol digit substitution online principal component 6 |
| Dur C1 | process durations principal component 1 |
| Dur C2 | process durations principal component 2 |
| Dur C4 | process durations principal component 4 |
| ECG C1 | ECG measures principal component 1 |
| ECG C2 | ECG measures principal component 2 |
| ECG C3 | ecg measures principal component 3 |
| ECG C6 | ECG measures principal component 6 |

| | |
|---|---|
| Family C1 | family history (parent's age) principal component 1 |
| Fluid Int. | fluid intelligence |
| Genetic C1 | genetic principal components and heterozygosity principal component 1 |
| Grip C1 | hand grip strength principal component 1 |
| Hearing | hearing signal-to-noise-ratio (snr) of triplet (left) |
| Illness C1 | non-cancer illness principal component 1 |
| Illness C4 | non-cancer illness principal component 4 |
| Loc C1 | location principal component 1 |
| Match | pairs matching |
| Match-o | pairs matching online |
| Matrix C1 | matrix pattern completion principal component 1 |
| Matrix C2 | matrix pattern completion principal component 2 |
| Matrix C3 | matrix pattern completion principal component 3 |
| Neuro | neuroticism score |
| ProMem C1 | prospective memory principal component 1 |
| RT C1 | reaction time principal component 1 |
| Sex | sex |
| Sex G C1 | genotype sex inference principal component 1 |
| Sex G C2 | genotype sex inference principal component 2 |
| Sleep | sleep duration per day |
| Smoke C1 | smoke principal component 1 |
| Time drive | time spent driving per day |
| Time TV | time spent watching television (tv) per day |
| Time walk | number of days walked 10+ minutes per week |
| Tower C1 | tower rearranging principal component 1 |
| Trail C1 | trail making principal component 1 |
| Trail-o C1 | trail making online principal component 1 |
| Trail-o C4 | trail making online principal component 4 |
| Travel | frequency of travelling from home to job workplace per week |
| Urine C1 | urine assays principal component 1 |
| Work | weekly length of working hour for main job |

**Table S2.** Dictionary of 67 phenotypes in the UK Biobank.

| ABCD field | Description |
|---|---|
| cbcl_scr_syn_anxdep_r | AnxDep CBCL Syndrome Scale (raw score) |
| cbcl_scr_syn_withdep_r | WithDep CBCL Syndrome Scale (raw score) |
| cbcl_scr_syn_somatic_r | Somatic CBCL Syndrome Scale (raw score) |
| cbcl_scr_syn_social_r | Social CBCL Syndrome Scale (raw score) |
| cbcl_scr_syn_thought_r | Thought CBCL Syndrome Scale (raw score) |
| cbcl_scr_syn_attention_r | Attention CBCL Syndrome Scale (raw score) |
| cbcl_scr_syn_rulebreak_r | RuleBreak CBCL Syndrome Scale (raw score) |
| cbcl_scr_syn_aggressive_r | Aggressive CBCL Syndrome Scale (raw score) |
| nihtbx_picvocab_uncorrected | NIH Toolbox Picture Vocabulary Test Age 3+ v2.0 Uncorrected Standard Score |
| nihtbx_flanker_uncorrected | NIH Toolbox Flanker Inhibitory Control and Attention Test Ages 8-11 v2.0 Uncorrected Standard Score |
| nihtbx_list_uncorrected | NIH Toolbox List Sorting Working Memory Test Age 7+ v2.0 Uncorrected Standard Score |
| nihtbx_cardsort_uncorrected | NIH Toolbox Dimensional Change Card Sort Test Ages 8-11 v2.0 Uncorrected Standard Score |
| nihtbx_pattern_uncorrected | NIH Toolbox Pattern Comparison Processing Speed Test Age 7+ v2.0 Uncorrected Standard Score |
| nihtbx_picture_uncorrected | NIH Toolbox Picture Sequence Memory Test Age 8+ Form A v2.0 Uncorrected Standard Score |
| nihtbx_reading_uncorrected | NIH Toolbox Oral Reading Recognition Test Age 3+ v2.0 Uncorrected Standard Score |
| nihtbx_fluidcomp_uncorrected | Cognition Fluid Composite Uncorrected Standard Score |
| nihtbx_cryst_uncorrected | Crystallized Composite Uncorrected Standard Score |
| nihtbx_totalcomp_uncorrected | Cognition Total Composite Score Uncorrected Standard Score |
| upps_y_ss_negative_urgency | UPPS-P for Children Short Form, Negative Urgency |
| upps_y_ss_lack_of_planning | UPPS-P for Children Short Form, Lack of Planning |
| upps_y_ss_sensation_seeking | UPPS-P for Children Short Form, Sensation Seeking |
| upps_y_ss_positive_urgency | UPPS-P for Children Short Form, Positive Urgency |
| upps_y_ss_lack_of_perseverance | UPPS: Lack of Perseverance (GSSF) |
| bis_y_ss_bis_sum | BIS/BAS: BIS Sum |
| bis_y_ss_bas_rr | BIS/BAS: BAS Reward Responsiveness |
| bis_y_ss_bas_drive | BIS/BAS: BAS drive |
| bis_y_ss_bas_fs | BIS/BAS: BAS Fun Seeking |
| pps_y_ss_number | Prodromal Psychosis Scale: Number of Yes Responses Sum |
| pps_y_ss_severity_score | Prodromal Psychosis: Severity Score Sum |
| pgbi_p_ss_score | Parent General Behavior Inventory SUM |
| pea_ravlt_sd_trial_vi_tc | RAVLT Short Delay Trial VI Total Correct |

| | |
|---|---|
| pea_ravlt_ld_trial_vii_tc | RAVLT Long Delay Trial VII Total Correct |
| pea_wiscv_trs | WISC-V Matrix Reasoning Total Raw Score |
| lmt_scr_perc_correct | Percentage correct of all 32 presented trials |
| lmt_scr_rt_correct | Average reaction for correct trials |
| lmt_scr_efficiency | LMT Efficiency |

**Table S3.** Dictionary of 36 phenotypes in the ABCD study.

| GSP field | Description |
|---|---|
| Flanker_S_CORRpc | Flanker Accuracy |
| Average of MenRot_80_CORRpc, MenRot_120_CORRpc, and MentRot_160_CORRpc | Mental Rotation Accuracy |
| Shipley_Vocab_Raw | Shipley Vocabulary |
| Matrix_WAIS | WAIS – Matrix Reasoning |
| STAI_tAnxiety | Speilberger Trait Anxiety |
| STAI_sAnxiety | Speilberger State Anxiety |
| POMS_TotMdDisturb | Total Mood Disturbance |
| Barratt_tot | Barratt Impulsivity |
| MindWandering_Freq | Mind Wandering Frequency |
| NEO_N | NEO – Neuroticism |
| NEO_E | NEO – Extraversion |
| NEO_O | NEO – Openness to Experience |
| NEO_A | NEO – Agreeableness |
| NEO_C | NEO – Conscientiousness |
| TCI_Novelty | Novelty Seeking |
| TCI_RewardDependance | Reward Dependence |
| TCI_HarmAvoidance | Harm Avoidance |
| DOSPERT_taking | Risk Raking |
| DOSPERT_perception | Risk Perception |
| BISBAS_BAS_Drive | Behavioral Activation – Drive |
| BISBAS_BAS_Fun | Behavioral Activation – Fun |
| BISBAS_BAS_Reward | Behavioral Activation – Reward |
| BISBAS_BIS | Behavioral Inhibition |

**Table S4.** Dictionary of 23 phenotypes in the GSP dataset.

| HBN field | Description |
|---|---|
| SDQ_Conduct_Problems | SDQ: Conduct Problems Scale |
| SDQ_Difficulties_Total | SDQ: Total Difficulties Score |
| SDQ_Emotional_Problems | SDQ: Emotional Problems Scale |
| SDQ_Externalizing | SDQ: Externalizing Score |
| SDQ_Generating_Impact | SDQ: Generating Impact Scores |
| SDQ_Hyperactivity | SDQ: Hyperactivity Scale |
| SDQ_Internalizing | SDQ: Internalizing Score |
| SDQ_Peer_Problems | SDQ: Peer Problems Scale |
| SDQ_Prosocial | SDQ: Prosocial Scale |
| SRS_AWR_T | SRS: Social Awareness T-Score |
| SRS_COG_T | SRS: Social Cognition T-Score |
| SRS_COM_T | SRS: Social Communication T-Score |
| SRS_DSMRRB_T | SRS: Restricted Interests and Repetitive Behavior T-Score |
| SRS_MOT_T | SRS: Social Motivation T-Score |
| SRS_RRB_T | SRS: Restricted Interests and Repetitive Behavior T-Score |
| SRS_SCI_T | SRS: Social Communication and Interaction T-Score |
| SCQ_Total | Social Communication Questionnaire Total Score |
| ASSQ_Total | Autism Spectrum Screening Questionnaire Total Score |
| SWAN_IN_Avg | SWAN Rating Scale for ADHD: Inattention Average |
| SWAN_HY_Avg | SWAN Rating Scale for ADHD: Hyperactivity Average |
| ARI_S_Total_Score | Affective Reactivity Index Self-Report Total Score |
| ARI_P_Total_Score | Affective Reactivity Index Parent-Report Total Score |
| CBCL_AD_T | CBCL: Anxious/Depressed T Score |
| CBCL_WD_T | CBCL: Withdrawn/Depressed T Score |
| CBCL_SC_T | CBCL: Somatic Complaints T Score |
| CBCL_SP_T | CBCL: Social Problems T Score |
| CBCL_TP_T | CBCL: Thought Problems T Score |
| CBCL_AP_T | CBCL: Attention Problems T Score |
| CBCL_RBB_T | CBCL: Rule Breaking Behavior T Score |
| CBCL_AB_T | CBCL: Aggressive Behavior T Score |
| CBCL_OP | CBCL: Other Problems Raw Score |
| CBCL_Int_T | CBCL: Internalizing T Score |
| CBCL_Ext_T | CBCL: Externalizing T Score |
| CELF_Total | Clinical Evaluation of Language Fundamentals Total Score |
| WIAT_Num_Stnd | WIAT: Numerical Operations Standard Score |

| | |
|---|---|
| WIAT_Pseudo_Stnd | WIAT: Pseudo-word Decoding Standard Score |
| WIAT_Spell_Stnd | WIAT: Spelling Standard Score |
| WIAT_Word_Stnd | WIAT: Word Reading Standard Score |
| NIH7_Card | Dimensional Change Card Sort Age 3+ Age-adjusted Scale Score |
| NIH7_Flanker | Flanker Inhibitory Control and Attention Age 3+ Age-adjusted Score |
| NIH7_List | List Sorting Working Memory Age 7+ Age-adjusted Score |
| NIH7_Pattern | Pattern Comparison Process Speed 7+ Age-adjusted Score |

**Table S5.** Dictionary of 42 phenotypes in the HBN dataset.

| eNKI-RS field | Description |
|---|---|
| ANT_01 | Alert Effect |
| ANT_02 | Orienting Effect |
| ANT_03 | Conflict Effect |
| DF_23 | Total Set Loss Designs Scaled |
| DF_25 | Total Repeated Designs Scaled |
| DF_27 | Total Attempted Designs Scaled |
| DF_29 | Percent Design Accuracy Scaled Score |
| DKEFSTMT_18 | Visual Scanning Scaled |
| DKEFSTMT_19 | Number Sequencing Scaled |
| DKEFSTMT_20 | Letter Sequencing Scaled |
| DKEFSTMT_21 | Number-Letter Switching Scaled |
| DKEFSTMT_22 | Motor Speed Scaled |
| DKEFSTMT_24 | Combined Number + Letter Sequencing Composite |
| DKEFSTMT_26 | Switching vs Visual Scanning Scaled |
| DKEFSTMT_28 | Switching Vs Number Sequencing Scaled |
| DKEFSTMT_30 | Switching Vs Letter Sequencing Scaled |
| DKEFSTMT_32 | Switching vs combined Number + letter sequencing Scaled |
| DKEFSTMT_34 | Switching Vs Motor Speed Scaled |
| TOWER_47 | Total Achievement Score Scaled |
| TOWER_49 | Total Rule Violation Scaled |
| TOWER_51 | Mean First Move Time Scaled |
| TOWER_53 | Time Per Move Ratio Scaled |
| TOWER_55 | Rule Violations Per Item Ratio Scaled |
| TOWER_57 | Move Accuracy Ratio Scaled |
| DEHQ_16 | Laterality Index |
| PTSDCH_55 | PTSD Severity: Overall Score |

| | |
|---|---|
| INT_12 | VCI Composite Score |
| INT_13 | PRI Composite Score |
| INT_14 | FSIQ - 4 Composite Score |
| WIAT_04 | Word Reading Subtest Standard Score |
| WIAT_05 | Numerical Operations Subtest Standard Score |
| WIAT_06 | Spelling Subtest Standard Score |
| WIAT_08 | Composite Standard Score |
| PENNCNP_0131 | ER40 Correct Responses Median Response Time |
| PENNCNP_0136 | ER40 Correct Anger Identifications |
| PENNCNP_0137 | ER40 Correct Fear Identifications |
| PENNCNP_0139 | ER40 Correct Happy Identifications |
| PENNCNP_0140 | ER40 Correct No Emotion Identifications |
| VF_32 | Category Fluency Scaled |
| VF_34 | Category Switching Scaled |
| VF_35 | Category Switching: Total Switching Accuracy |
| VF_37 | Letter Fluency Vs Category Fluency Scaled |
| VF_39 | Category Switching Vs Category Fluency Scaled |
| VF_41 | First Interval Total Correct Scaled |
| VF_43 | Second Interval Correct Scaled |
| VF_45 | Third Interval Correct Scaled |
| VF_47 | Fourth Interval Correct Scaled |
| VF_49 | Set Loss Errors Scaled |
| VF_51 | Repetition Errors Scaled |
| VF_54 | Percent Set Loss Errors Scaled |
| DKEFSCWI_13 | Color Naming Scaled |
| DKEFSCWI_14 | Word Reading Scaled |
| DKEFSCWI_15 | Inhibition Scaled |
| DKEFSCWI_16 | Inhibition/Switching Scaled |
| DKEFSCWI_18 | Combined Naming + Reading Comp Scaled |
| DKEFSCWI_20 | Inhibition vs color naming Scaled |
| DKEFSCWI_22 | Inhibition Vs Combined Naming + reading Scaled |
| DKEFSCWI_24 | Inhibition Switching Vs Inhibition Scaled |
| DKEFSCWI_26 | Inhibition/Switching Vs Color Naming Scaled |
| DKEFSCWI_28 | Inhibition/Switching Vs Word Reading Scaled |
| DKEFSCWI_40 | Total Errors Inhibition/Switching Scaled Score |

**Table S6.** Dictionary of 61 phenotypes in the eNKI-RS dataset.

| HCP-YA field | Description |
|---|---|
| PicSeq_Unadj | Visual Episodic Memory |
| CardSort_Unadj | Cognitive Flexibility (DCCS) |
| Flanker_Unadj | Inhibition (Flanker Task) |
| PMAT24_A_CR | Fluid Intelligence (PMAT) |
| ReadEng_Unadj | Vocabulary (Pronunciation) |
| PicVocab_Unadj | Vocabulary (Picture Matching) |
| ProcSpeed_Unadj | Processing Speed |
| DDic_AUC_40K | Delay Discounting |
| VSPLOT_TC | Spatial Orientation |
| SCPT_SPEC | Sustained Attention – Spec. |
| ListSort_Unadj | Working Memory (List Sorting) |
| MMSE_Score | Cognitive Status (MMSE) |
| PSQI_Score | Sleep Quality (PSQI) |
| Endurance_Unadj | Walking Endurance |
| GaitSpeed_Unadj | Walking Speed |
| Dexterity_Unadj | Manual Dexterity |
| Strength_Unadj | Grip Strength |
| Taste_Unadj | Taste Intensity |
| Emotion_Task_Face_Acc | Emotional Face Matching |
| Language_Task_Math_Avg_Difficulty_Level | Arithmetic |
| Language_Task_Story_Avg_Difficulty_Level | Story Comprehension |
| Relational_Task_Acc | Relational Processing |
| WM_Task_Acc | Working Memory (N-back) |
| NEOFAC_A | Agreeableness (NEO) |
| NEOFAC_O | Openness (NEO) |
| NEOFAC_C | Conscientiousness (NEO) |
| NEOFAC_E | Extraversion (NEO) |
| AngAggr_Unadj | Anger – Aggression |
| FearAffect_Unadj | Fear – Affect |
| Sadness_Unadj | Sadness |
| LifeSatisf_Unadj | Life Satisfaction |
| MeanPurp_Unadj | Meaning & Purpose |
| Loneliness_Unadj | Loneliness |
| PercStress_Unadj | Perceived Stress |
| SelfEff_Unadj | Self-Efficacy |

**Table S7.** Dictionary of 35 phenotypes in the HCP-YA dataset.

| HCP-Aging field | Description |
|---|---|
| LifeSatisf | Life Satisfaction |
| MeanPurp | Meaning & Purpose |
| PosAffect | Positive Affect |
| Sadness | Sadness |
| FearAffect | Fear – Affect |
| FearSomat | Fear – Somatic |
| AngAffect | Anger – Affect |
| AngHostil | Anger – Hostility |
| AngAggr | Anger – Aggression |
| ER40ANG | ER40 Correct Anger Identifications |
| ER40FEAR | ER40 Correct Fear Identifications |
| ER40NOE | ER40 Correct No Emotion Identifications |
| ER40SAD | ER40 Correct Sad Identifications |
| EmotSupp | Emotion Support |
| InstruSupp | Instrumental Support |
| Friendship | Friendship |
| Loneliness | Loneliness |
| PercReject | Perceived Rejection |
| PercHostil | Perceived Hostility |
| PercStress | Perceived Stress |
| SelfEff | Self-Efficacy |
| NEOFAC_A | Agreeableness (NEO) |
| NEOFAC_O | Openness (NEO) |
| NEOFAC_C | Conscientiousness (NEO) |
| NEOFAC_N | Neuroticism (NEO) |
| NEOFAC_E | Extraversion (NEO) |
| Endurance | Walking Endurance |
| GaitSpeed | Walking Speed |
| Strength | Grip Strength |
| PicSeq | Visual Episodic Memory |
| CardSort | Cognitive Flexibility (DCCS) |
| Flanker | Inhibition (Flanker Task) |
| ReadEng | Vocabulary (Pronunciation) |
| PicVocab | Vocabulary (Picture Matching) |
| ProcSpeed | Processing Speed |
| DDisc_AUC_40K | Delay Discounting |

| | |
|---|---|
| ListSort | Working Memory (List Sorting) |
| raw_vat | Raw Score for the Visual Acuity |
| raw_pain | Raw Score for the Pain Interference |
| winright_ncorr | Words in Noise Number Correct Right |
| psqi_total | Sleep Quality (PSQI) |
| tmta_raw | Trailmaking test part A: Time to completion |
| tmtb_raw | Trailmaking test Part B: Time to completion |
| pea_ravlt_sd_tc | RAVLT Short Delay Total Correct |
| moca_total | Montreal Cognitive Assessment Total Score |

**Table S8.** Dictionary of 45 phenotypes in the HCP-Aging dataset.

*S3. Bootstrapping procedure*

Following our previous study (He et al., 2022), performance difference between algorithms were evaluated using a bootstrapping approach. More specifically, we performed bootstrap sampling of K-shot 1,000 times, i.e., we randomly picked K participants repeatedly with replacements from the meta-test dataset (so there might be duplicated participants). For each of the 1,000 bootstrapped samples, we applied all 5 algorithms (classical KRR, transfer learning, meta-matching with stacking, meta-matching with dataset stacking, and multilayer meta-matching), yielding prediction accuracies (Pearson's correlation and COD) in the remaining participants in the meta-test dataset. Note that the remaining participants might be more than N – K participants because some of the K participants might be duplicated (because of sampling with replacement).

Therefore, for each algorithm, we had 1,000 bootstrapped Pearson's correlation and 1,000 bootstrapped COD values. To explain the statistical testing procedure, we focus on COD in the following explanation for clarity, since the procedure for Pearson's correlation was the same as that for COD.

To compute the statistical difference between algorithm A and algorithm B, we calculated the difference between 1,000 bootstrapped samples for COD from algorithm A and 1,000 bootstrapped samples for COD from algorithm B, and we fitted a Gaussian distribution to the difference of 1,000 COD between A and B. We denoted the mean, standard deviation, and cumulative distribution function of the Gaussian distribution as $\mu$, $\sigma$, and $\mathrm{CDF}_{\mathrm{A\text{-}B}}$, respectively. We then computed p-value by comparing 0 with $\mathrm{CDF}_{\mathrm{A\text{-}B}}$, i.e.:

$$p = \begin{cases} 2 \times \mathrm{CDF}_{\mathrm{A-B}}(0) & \text{, if } \mu \geq 0 \\ 2 \times \left(1 - \mathrm{CDF}_{\mathrm{A-B}}(0)\right) & \text{, if } \mu < 0 \end{cases}$$

P values were computed between all pairs of algorithms. Since there were multiple comparisons for different values of K (K = 10, 20, 50, 100, 200), different accuracy metrics (Pearson's correlation and COD), and for different pairs of algorithms, multiple comparisons were corrected using a false discovery rate (FDR) of $q < 0.05$. FDR was applied to all K-shots and across all pairs of algorithms and both evaluation metrics.

### Supplementary Results

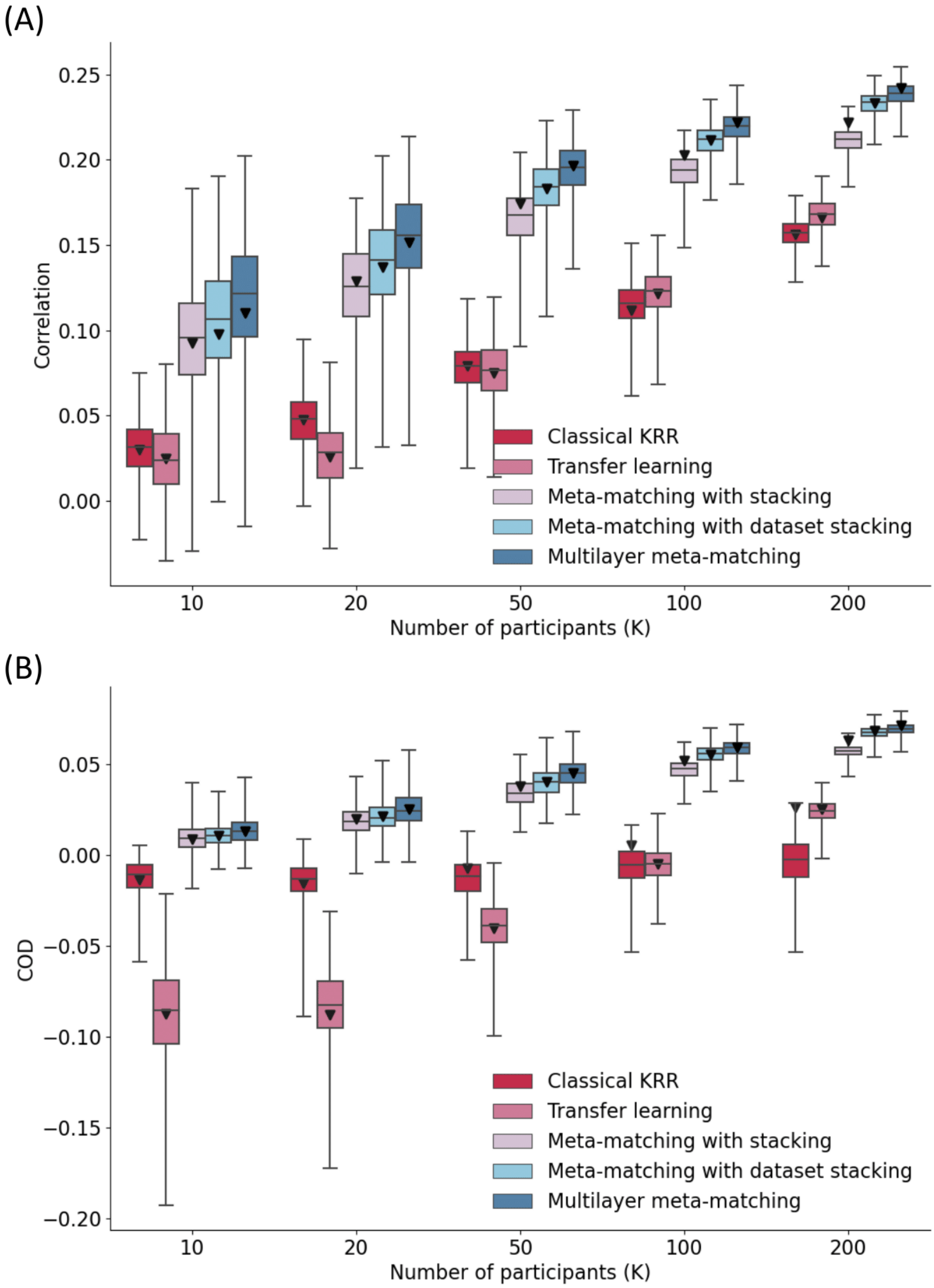


**Figure S1. Multilayer meta-matching outperformed meta-matching with stacking on HCP-YA bootstrap experiment.** (A) Phenotypic prediction performance in terms of Pearson's correlation (averaged across 35 meta-test phenotypes) on HCP-YA bootstrap samples. Horizontal axis is the number of few-shot participants (K); Vertical axis is Pearson's correlation of phenotypic prediction. Boxplots represent variability across 1,000 bootstrapped K-shot, and different colors stand for different approaches. Inverted triangles represent the average prediction performance (correlation) on 100 random K-shot. (B) Prediction performance in terms of COD on HCP-YA bootstrap samples. All settings were the same as (A) but used COD as the metric.

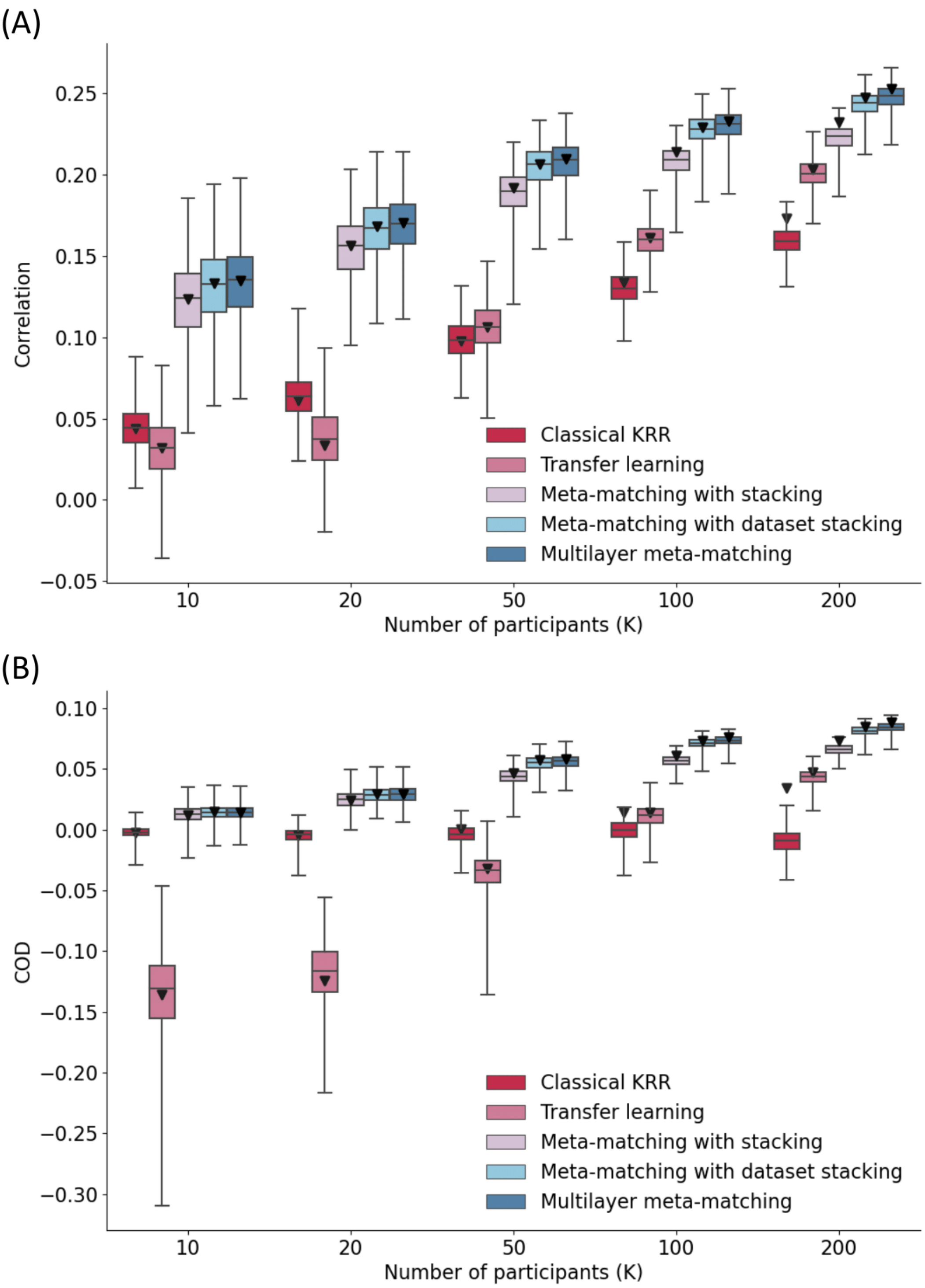


**Figure S2. Multilayer meta-matching outperformed meta-matching with stacking on HCP-Aging bootstrap experiment.** (A) Phenotypic prediction performance in terms of Pearson's correlation (averaged across 45 meta-test phenotypes) on HCP-Aging bootstrap samples. Horizontal axis is the number of participants (K); Vertical axis is Pearson's correlation of phenotypic prediction. Boxplots represent variability across 1,000 bootstrapped K-shot, and different colors stand for different algorithms. Inverted triangles represent the average prediction performance (correlation) on 100 random K-shot. (B) Prediction performance in terms of COD on HCP-Aging bootstrap samples. All settings were the same as A but used COD as the metric.
